# Control of antigen presentation on MHC-I by a bacterial secretion system

**DOI:** 10.1101/2025.11.19.689332

**Authors:** Owen Leddy, Rachel McGinn, Benjamin Allsup, Madison Leone, Penny Timms, Matias Gutierrez-Gonzalez, Ahmed S. Fahad, Ryan Milligan, Wilfredo F. Garcia-Beltran, Brandon J. DeKosky, Forest M. White, Bryan D. Bryson

**Author notes:** Equal contribution (Owen Leddy, Rachel McGinn). Equal contribution (Forest M. White, Bryan D. Bryson).

## Abstract

*Mycobacterium tuberculosis* (Mtb) remains one of the world’s leading infectious killers. Although CD8□ T cells contribute to immune control of tuberculosis, the pathways through which bacterial antigens access major histocompatibility complex class I (MHC-I) antigen presentation remain incompletely defined. Here, we show that the activity of an Mtb secretion system actively promotes antigen presentation on MHC-I. Using quantitative immunopeptidomics, host and bacterial genetic perturbations, and T cell activation assays, we demonstrate that presentation of Mtb-derived peptides on MHC-I requires the ESX-1 type VII secretion system. Presentation of these peptides proceeds in a manner dependent on the transporter associated with antigen processing (TAP) but independent of host cell mechanisms such as autophagy or MPEG1-mediated pore formation. Chemical induction of phagosomal membrane damage fails to restore antigen presentation in the absence of ESX-1 activity, suggesting that pathogen-encoded secretion, not nonspecific membrane rupture, governs access to MHC-I antigen processing pathways. These findings reveal a secretion system–driven mechanism of antigen presentation, redefining how mycobacteria interface with host MHC-I pathways, potentially informing tuberculosis vaccine design strategies, and highlighting a potential route for synthetic antigen delivery to the cytosol in therapeutics and vaccination.

**One-sentence summary:** Presentation of *Mycobacterium tuberculosis* antigens on MHC class I through a cytosolic pathway depends on a bacterial secretion system rather than host response and cross-presentation pathways.

## Introduction

Cell-mediated immunity is an essential component of the immune response to cancer and infection. Defining the mechanisms by which cell-mediated immunity can be effectively elicited and harnessed can thus enhance and accelerate the development of next generation immunotherapies and vaccines. Harnessing CD8+ T cells is of great interest given their ability to potently drive clearance of the target cell and the pathogen in the case of pathogen-infected cells. Understanding how antigens are seen by CD8+ T cells in the context of infection and cancer can therefore provide novel molecular strategies to enhance immunosurveillance. A major gap in the field to date has been defining the molecular mechanisms by which antigens that are localized in endomembrane compartments are seen by the cytosolic machinery required for antigen presentation on MHC-I. Several recent studies have provided novel insights into the mechanism by which antigens are cross-presented from endomembrane-containing compartments to the cytosol; however, these studies largely focused on host factors that regulate these processes.^1–3^

Antigen presentation on major histocompatibility complex molecules is central to effective induction of cell-mediated immunity in infection, cancer, and other diseases. T cell-mediated immunity is a cornerstone of the immune response to *Mycobacterium tuberculosis* (Mtb), and evidence from animal models suggests that CD8+ T cells can contribute to protective immunity against tuberculosis (TB). This implies that optimal design of TB vaccines will require leveraging immunity mediated by CD8+ as well as CD4+ T cells. Non-human primates that were infected with Mtb, cured, and re-challenged showed increased bacterial burden if CD8+ T cells were depleted prior to re-challenge.^4^ Mice deficient in MHC-I also succumb more rapidly to Mtb infection.^5^ However, *ex vivo* CD8+ T cells from Mtb-infected mice^6^ and from human participants in a phase I clinical trial of a virus-vectored candidate TB vaccine^7^ inefficiently recognized Mtb-infected phagocytes, suggesting that not all Mtb-specific CD8+ T cells can contribute to immune control due to a lack of cognate antigen in the MHC-I repertoire of infected cells. Therefore, understanding the mechanisms by which Mtb peptides are processed and presented on MHC-I by infected cells may help improve future TB vaccine designs.

In order to prime CD8+ T cells to directly recognize infected cells, vaccines against Mtb infection must drive presentation of the same peptide epitopes that are presented on MHC-I during infection. Whereas viral antigens are produced by the host cell translational machinery and therefore are exposed to the same antigen processing and presentation pathways as endogenous proteins, antigens derived from bacterial or eukaryotic pathogens may not all be accessible to host antigen processing pathways if they are not exposed outside the microbial cell. For pathogens that reside in a membrane-bound intracellular compartment, this compartment may represent an additional barrier restricting which pathogen-derived proteins are accessible to host antigen processing pathways, resulting in a limited repertoire of MHC-I epitopes in infected cells.^8^ Understanding the mechanisms by which antigens derived from a phagosome-resident pathogen like Mtb access antigen processing pathways in infected macrophages may therefore help inform the selection of appropriate vaccine antigens and delivery strategies.

Antigens derived from phagocytosed cargo are typically thought to be presented on MHC-I through two broad categories of cross-presentation pathways: 1) translocation to the cytosol by host pore-forming proteins, where they can be processed and presented through the same pathways as endogenous MHC-I antigens, or 2) processing and loading on MHC-I within the phagosome itself.^3,9–12^ A third possibility specific to pathogens that survive and remain active in the phagosome is that the pathogen itself may enable antigens to access the host cytosol by disrupting the phagosome membrane^13^ or selectively injecting effector proteins into the host cytosol using specialized secretion systems.^14^

In prior work, we used untargeted mass spectrometry (MS) analysis of the MHC-I repertoire of Mtb-infected human macrophages to identify antigenic targets that could potentially be recognized by CD8+ T cells.^15^ We found that Mtb peptides presented on MHC-I were predominantly derived from substrates of Mtb’s type VII secretion systems (T7SSs) – molecular machines that export Mtb proteins across the cell envelope and have roles in virulence,^16^ nutrient acquisition,^17^ and outer membrane biogenesis.^18^ We identified MHC-I peptides derived from substrates of three distinct T7SSs (ESX-1, ESX-3, and ESX-5). These results suggested that T7SS substrates preferentially gain access to MHC-I antigen processing pathways relative to other Mtb proteins. We and others have also identified the activity of the ESX-1 T7SS as a requirement for efficient Mtb antigen presentation on MHC-I.^15,19^ ESX-1 mediates rupture of the phagosome membrane enclosing intracellular Mtb,^20,21^ and inhibitors of components of cytosolic antigen processing pathways have previously been shown to inhibit CD8+ T cell recognition of Mtb infected cells.^22^ Here, we used a combination of quantitative targeted MS, host and bacterial genetic perturbations and additional systems biology techniques to demonstrate that the Mtb ESX-1 secretion system is required for efficient presentation of Mtb-derived antigens on MHC-I rather than through cross-presentation pathways driven solely by the host cell. These data expand the potential molecular strategies that can be harnessed to effectively elicit CD8+ T cell immunity.

## Results

### The ESX-1 secretion system of Mtb is required for recognition by Mtb-specific CD8+ T cells

In studying presentation of Mtb-derived peptides on MHC-I, we demonstrated using mass spectrometry that loss of activity of Mtb’s ESX-1 type VII secretion system disrupts presentation of Mtb-derived peptides on MHC-I.^15^ These data suggested that there may be pathogen-mediated mechanisms that control presentation of Mtb-derived peptides on MHC-I. We first sought to validate the results of our previous study using a T cell reporter line specific for an Mtb-derived peptide whose sequence is shared across the ESX-5 substrates EsxJ, EsxK, EsxP, and EsxW (hereinafter referred to as EsxJKPW_24-34_). Lewinsohn and colleagues had independently identified CD8+ T cells from individuals with evidence of prior Mtb infection capable of recognizing peptides encoded by the Mtb genome. Among the T cells that recognized Mtb-derived peptides, they identified a T cell clone at a circulating frequency of ∼0.03% that recognized the exact epitope we identified by MS restricted by the same HLA allotype,^23^ providing evidence of the *in vivo* relevance of our studies of Mtb-derived antigen presentation in cells infected *in vitro*. We cloned the T cell receptor from this T cell clone into SKW3 reporter cells and confirmed using HLA-matched human monocyte-derived macrophages (hMDMs) that loss of ESX-1 activity disrupts presentation of EsxJKPW_24-34_ (Supplementary Figure 1). Together with our prior MS data, these T cell coculture results support a model in which the activity of a pathogen’s secretion system controls antigen presentation by infected cells *in vitro*.

There are several models that could explain how the ESX-1 secretion system influences Mtb-derived antigen presentation. ESX-1 controls secretion of specific Mtb proteins like EsxA and EsxB,^24^ galectin and P62 puncta formation at the phagosome,^15,21,25,26^ and type 1 interferon induction (Figure 1A).^15,27–29^ We previously demonstrated that exogenous addition of type 1 interferons does not rescue presentation of Mtb-derived antigens.^13^ However, the contributions of other pathways towards ESX-1’s influence on Mtb antigen presentation on MHC-I remained unclear.

**Figure 1.**
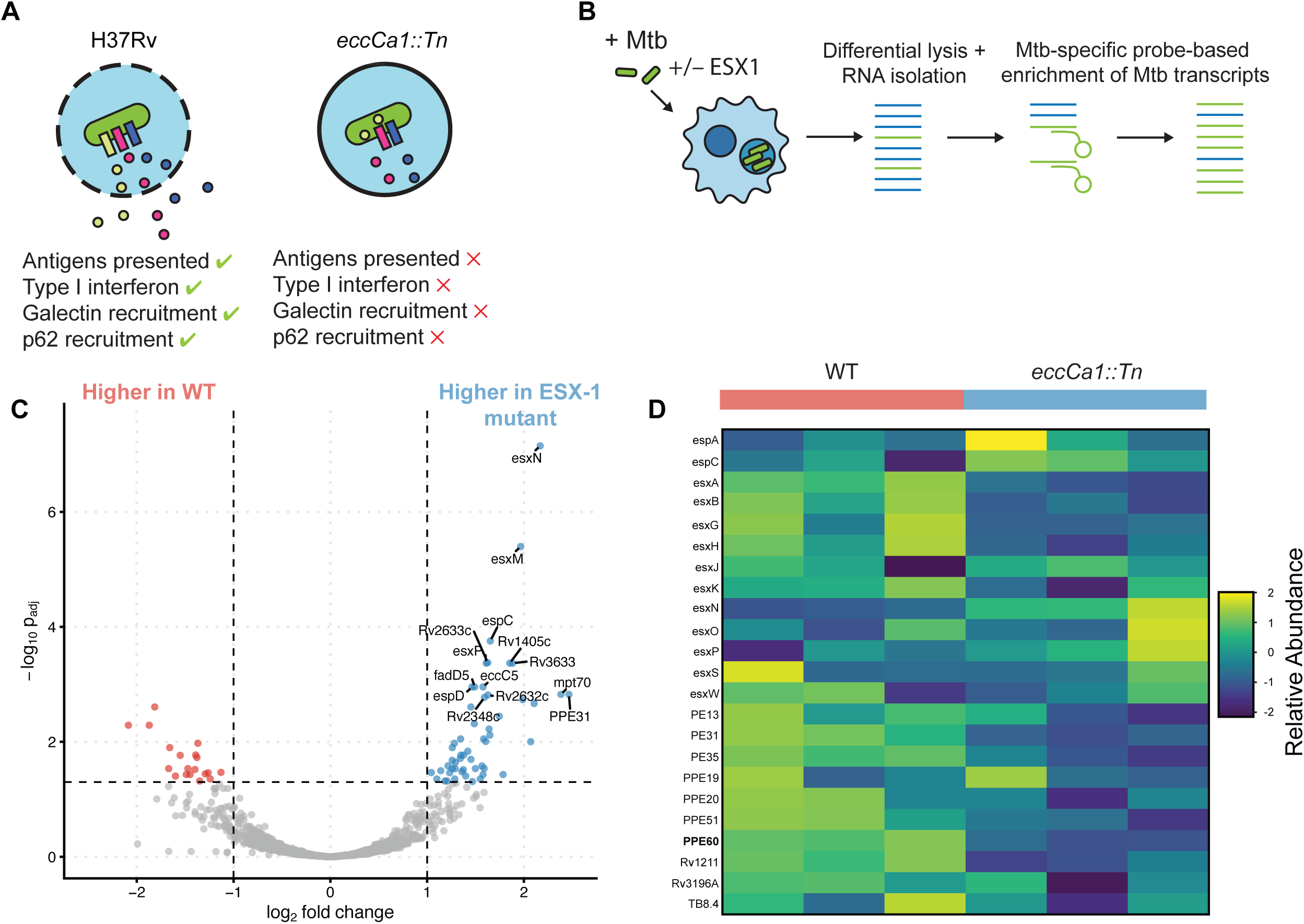
*Mtb* antigens presented on MHC-I are not transcriptionally downregulated in the absence of ESX-1 activity. A) Schematic representation of the effects of loss of ESX-1 function on *Mtb* interactions with the host cell. H37Rv denotes wild-type Mtb and *eccCa1::Tn* denotes ESX-1 loss of function mutant. B) Schematic overview of transcriptomics workflow for analysis of intracellular Mtb gene expression in hMDMs. C) Volcano plot of gene expression by intracellular Mtb comparing WT H37Rv and *eccCa1::Tn*. Differential gene expression was analyzed using DESeq2 with apeGLM-estimated log2 fold change shrinkage. Adjusted p-values were determined using a Wald test, using the Benjamini-Hochberg correction for multiple hypothesis testing. D) Heatmap of relative transcript abundance [row-wise z-score of log(transcripts per million + 1)] for Mtb genes encoding peptides presented on MHC-I identified in previous studies.^15,37^ Each column represents a biological replicate using hMDMs from a different donor.

### Loss of ESX-1 activity does not abrogate expression of genes encoding Mtb antigens presented on MHC-I

We sought to systematically test a range of models that might explain the observed reduction in Mtb-derived antigen presentation on MHC-I in the absence of ESX-1 activity. ESX-1 activity influences how Mtb-containing phagosomes mature, and it is possible that the environmental signals from mature phagolysosomes influence transcriptional regulation of Mtb antigens that are presented on MHC-I.^30^ To test whether the dependence of Mtb MHC-I antigen presentation on ESX-1 was the result of changes in antigen expression at the transcriptional level, we compared the gene expression profile of either wild-type (WT) Mtb H37Rv or an Mtb mutant lacking ESX-1 activity (*eccCa1::Tn*) at 24 hours post-infection in hMDMs, a time point when ESX-1-dependent antigen presentation is already detectable (Figure 1B). Given the small fraction of Mtb mRNAs in the pool of RNA from Mtb-infected hMDMs, we used a probe-based approach to enrich cDNAs derived from Mtb mRNAs. We measured differential gene expression of WT and ESX-1 mutant Mtb using DESeq2,^31^ and we applied Bayesian shrinking to log2 fold changes using the software package apeGLM^32^ to reduce noise in fold changes for genes with a low expression level (Figure 1C). We examined the expression of a curated list of Mtb genes encoding antigens we previously rigorously validated to be presented on MHC-I by Mtb-infected hMDMs or human monocyte-derived dendritic cells. Genes encoding MHC-I epitopes that we previously showed were ESX-1-dependent using quantitative MS (*esxG*, *esxS*, *esxA*, *esxB, esxJ, esxK, esxP, esxW*)^15^ were not differentially expressed between WT and ESX-1 mutant Mtb (Figure 1C-D, Supplementary Table 1). Of all the Mtb genes encoding MHC-I epitopes we had previously detected by immunopeptidomics, only *ppe60* was significantly downregulated in the ESX-1 mutant compared to WT (Figure 1C-D, Supplementary Table 1). In our immunopeptidomics data, the epitope detected from PPE60 is shared with PPE19,^15^ so we cannot attribute the peptide solely to either PPE19 or PPE60. *ppe19* was not significantly differentially expressed across the two experimental conditions. Taken together, these data argue that the dependence of Mtb MHC-I epitope presentation on ESX-1 activity cannot be attributed to differential gene expression at the transcriptional level.

### Mtb-derived antigens presented on MHC-I require TAP for presentation

A key role of ESX-1 during infection is disruption of the phagosomal membrane, leading to the recruitment of galectins to the phagosome, as well as other proteins associated with endomembrane compartment damage. We hypothesized that ESX-1 is required for Mtb antigen presentation on MHC-I because antigens must access the host cell cytosol in order to be presented. If access to the host cytosol is required for presentation of Mtb antigens on MHC-I, we would predict that the transporter associated with antigen processing (TAP) would be required to import cytosolically processed Mtb peptides into the endoplasmic reticulum (ER) for loading onto MHC-I^33^ (Figure 2A).

**Figure 2.**
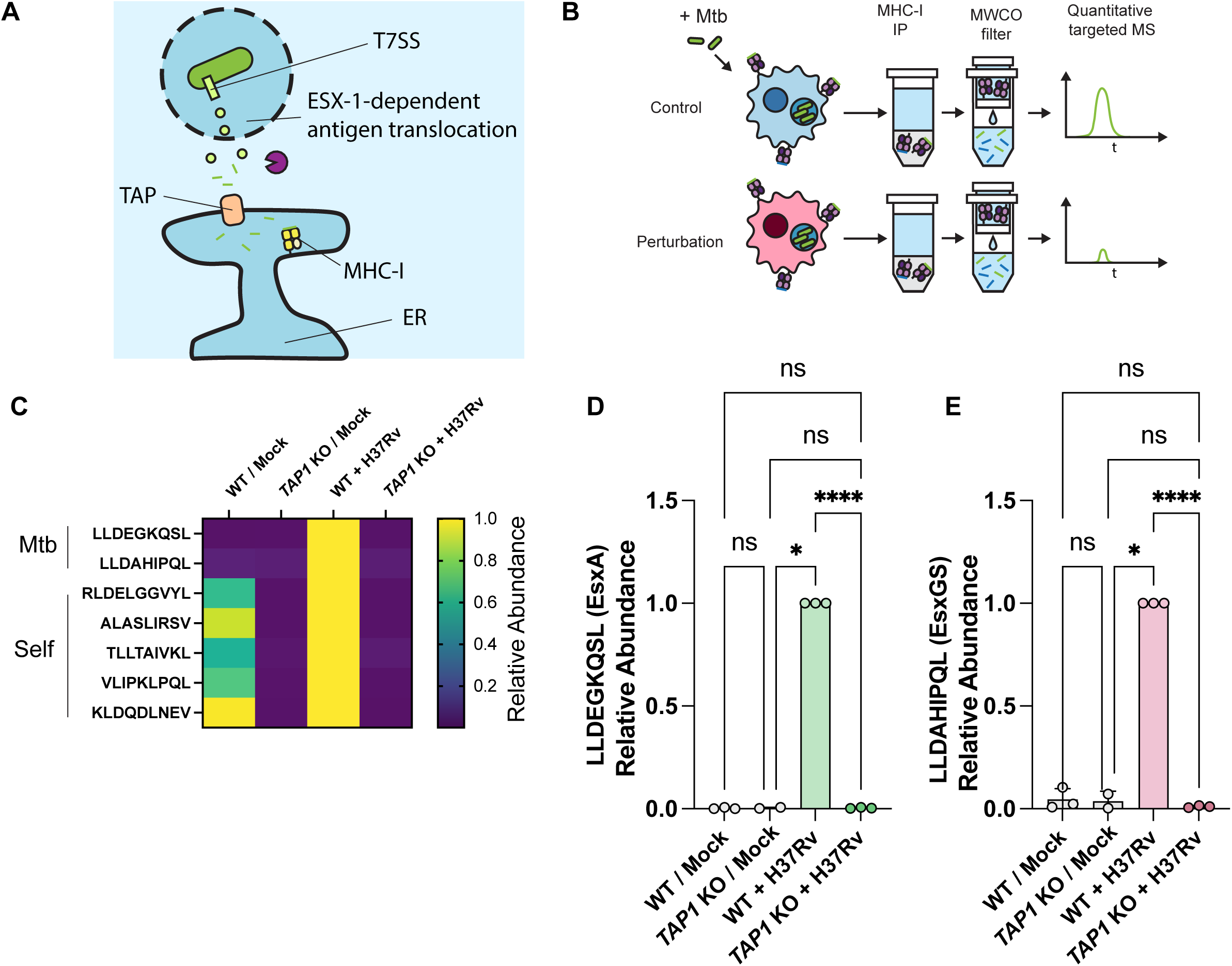
Presentation of *Mtb* antigens on MHC-I is TAP-dependent. A) Schematic illustration of a proposed model of *Mtb* MHC-I antigen presentation via a cytosolic pathway. B) Schematic illustration of the SureQuant targeted MS approach for quantifying *Mtb* MHC-I peptides. (MWCO: molecular weight cutoff.) (C-E) TAP1 knockout THP-1 macrophages or the parental wild-type line were infected with *Mtb* at MOI 2.5 and harvested for MHC-I peptide quantification by SureQuant 24 hours post-infection. C) Heatmap of relative abundances of MHC-I peptides (mean of n = 3 replicate experiments). D) Relative abundance of EsxA_28-36_ presented on MHC-I for n = 3 replicate experiments. E) Relative abundance of EsxGS_3-11_ presented on MHC-I for n = 3 replicate experiments.

To test this hypothesis, we infected THP-1 macrophages in which the TAP1 gene was knocked out, as well as the parental wild-type THP-1 line, and quantified presentation of two Mtb antigens presented on HLA-A*02:01 using the targeted MS method SureQuant MHC^34^ (Figure 2B). EsxA_28-36_ (sequence LLDEGKQSL) is derived from a substrate of the ESX-1 secretion system, while EsxGS_3-11_ (sequence LLDAHIPQL) is conserved between two paralogous substrates of the ESX-3 secretion system (EsxG and EsxS). These epitopes are both recognized by CD8+ T cells in HLA-A*02:01+ humans with prior Mtb exposure,^35,36^ and we previously detected both in the MHC-I repertoire of infected cells by MS.^15,37^ We also quantified five self MHC-I peptides as an internal control for overall MHC-I antigen presentation capacity. Knocking out TAP1 abolished presentation of all self-peptides and Mtb peptides on MHC-I, consistent with presentation of Mtb-derived peptides through a cytosolic antigen processing pathway (Figure 2C-E). These data showed that Mtb-derived MHC-I antigen presentation is TAP-dependent.

### Chemical modulation of phagosome membrane damage does not rescue presentation of Mtb-derived peptides on MHC-I

Our experiments thus far had shown that presentation of Mtb antigens on MHC-I is TAP-dependent, and our prior study showed that presentation was independent of type 1 interferon signaling.^15^ Therefore, we next sought to specifically examine a role for phagosome membrane damage in Mtb-derived antigen presentation on MHC-I. Given knowledge gaps in our understanding of the ESX-1 secretion system, very few tools outside of deletion or transposon mutants exist. To our knowledge, no mutations in Mtb ESX-1 components or substrates have been isolated that fully preserve secretion system activity but specifically abrogate phagosome membrane damage. For example, deletion of *esxA* abrogates recruitment of membrane damage markers to phagosomes,^21^ but also affects secretion of other ESX-1 substrates.^38,39^ Thus, testing the hypothesis that phagosome membrane damage alone is sufficient to enable *Mtb* MHC-I antigen presentation in the absence of other ESX-1 functions requires an orthogonal means of experimentally inducing phagosome membrane disruption. We therefore infected primary human monocyte-derived macrophages (hMDMs) with an *Mtb* strain lacking ESX-1 activity (*eccCa1::Tn*) and treated the cells with small molecules that have been previously shown to increase release of proteins residing in phagosomes into the cytosol.^40^ From this previous screen, we selected prazosin and gefitinib due to the magnitude of phagosomal antigen released in the previous screen. Like *Mtb*, intracellular *Mycobacterium marinum* co-localizes with membrane damage markers like galectins in an ESX-1-dependent manner, and treating cells infected with ESX-1-deficient *M. marinum* with prazosin has previously been shown to restore recruitment of these membrane damage markers.^41^ We first validated that treatment with prazosin or gefitinib restored galectin-3 colocalization with ESX-1 mutant Mtb. Whereas Mtb co-localized with the membrane damage marker galectin-3 in cells infected with wild-type Mtb, co-localization was greatly reduced in cells infected with ESX-1 mutant Mtb, as previously observed.^15,26^ Treating cells infected with ESX-1 mutant Mtb with prazosin or gefitinib restored co-localization of Mtb with galectin-3 to wild-type levels (Figure 3A-B).

**Figure 3.**
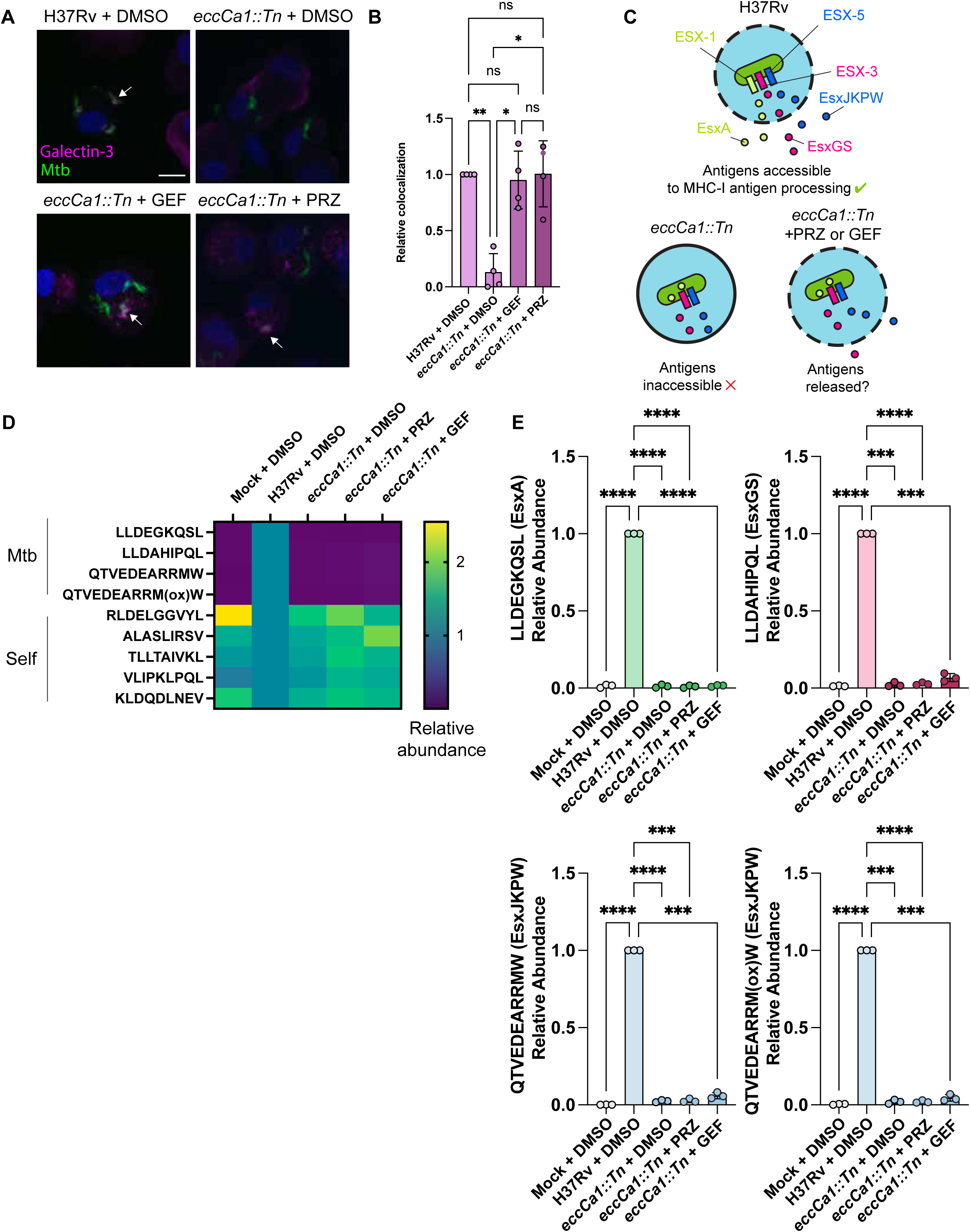
Drug-induced phagosome membrane disruption is not sufficient to rescue MHC-I antigen presentation in the absence of ESX-1 activity. (A-B) Primary hMDMs infected with wild-type or *eccCa1::Tn Mtb* at MOI 2.5 for 72 hours, treated with prazosin (PRZ), gefitinib (GEF), or DMSO alone for 2 hours, and stained for galectin 3. A) Representative fluorescence confocal microscope images. White arrows indicate intracellular bacteria co-localized with galectin 3. B) Relative quantification of colocalization between intracellular *Mtb* and galectin 3, as measured by analysis of local fluorescence intensity correlation (see Methods) for n = 4 donors. C) Schematic illustrating hypothesized mechanism of antigen release from the phagosome following membrane damage induced by ESX-1 or drug treatment. (D-E) Primary hMDMs expressing HLA-A*02:01 and HLA-B*57:01 were infected with wild-type or *eccCa1::Tn Mtb* at MOI 2.5 for 72 hours, treated with prazosin, gefitinib, or DMSO alone for 2 hours, and harvested for MHC-I antigen quantification by SureQuant. D) Heatmap of relative abundances of MHC-I peptides (mean of n = 3 donors). E) Relative abundance of EsxA_28-36_, EsxGS_3-11_, EsxJKPW_24-34_ (non-oxidized), and EsxJKPW_24-34_ (oxidized) presented on MHC-I for n = 3 donors.

We next sought to determine whether inducing galectin recruitment, putatively through membrane disruption, was sufficient to enable Mtb antigens to access cytosolic antigen processing pathways in the absence of ESX-1 activity. If this were the case, prazosin or gefitinib treatment of cells infected with ESX-1-deficient Mtb should rescue presentation of ESX-3 and ESX-5 substrates (secreted independently of ESX-1^42^) by releasing them into the cytosol from the phagosome lumen (Figure 3C). We infected hMDMs expressing HLA-A*02:01 and HLA-B*57:01 with WT or ESX-1 mutant Mtb and pulsed the cells with prazosin, gefitinib, or DMSO carrier alone for 2 hours, followed by lysis 6 hours later for MHC-I isolation. We used SureQuant MHC to quantify presentation of EsxA_28-36_ and EsxGS_3-11_ on HLA-A*02:01 and presentation of EsxJKPW_24-34_ (sequence QTVEDEARRMW, derived from a family of four ESX-5 substrates) on HLA-B*57:01. Like EsxA_28-36_ and EsxGS_3-11_, EsxJKPW_24-34_ is recognized by human CD8+ T cells^43^ and presented by infected cells.^15^ In agreement with our prior results^15^, presentation of EsxA_28-36_, EsxGS_3-11_ and EsxJKPW_24-34_ was greatly decreased in cells infected with ESX-1 mutant Mtb, relative to wild-type (Figure 3D-E). We found that treatment with prazosin or gefitinib was not sufficient to rescue presentation of Mtb antigens on MHC-I by cells infected with ESX-1 mutant Mtb (Figure 3D-E). Treating infected THP-1 macrophages with prazosin for 6 hours followed by lysis at 24 hours post-infection (mimicking the drug treatment timing used by Kozik et al. to demonstrate antigen release from phagosomes^40^) also did not rescue presentation of Mtb antigens, suggesting that this result was not merely due to suboptimal timing of drug treatment or cell harvesting (Supplementary Figure 2).

We and others previously showed that the dependence of Mtb MHC-I antigen presentation on ESX-1 activity was not solely due to lower bacterial burden in cells infected with ESX-1-deficient mutants relative to wild-type Mtb.^15,19^ We nonetheless wanted to determine whether intracellular Mtb was still present at comparable levels during infection with ESX-1-deficient Mtb. We therefore quantified GFP+ area (normalized by DAPI+ cell nuclei) in our fluorescence microscopy images, and found that while GFP+ area was modestly reduced on average in cells infected with ESX-1 mutant Mtb relative to wild-type (with or without treatment with prazosin or gefitinib), this <2-fold reduction was not sufficient to explain the much greater reduction in MHC-I antigen presentation (Supplementary Figure 3).

### Autophagy of intracellular Mtb is dispensable for Mtb antigen presentation on MHC-I

A plausible explanation for the failure of drug-induced phagosomal permeabilization to restore Mtb antigen presentation on MHC-I is that ESX-1 may trigger additional host responses that are necessary for effective MHC-I presentation. We therefore sought to identify other downstream consequences of ESX-1 activity that could contribute to antigen presentation and are not rescued by prazosin or gefitinib treatment.

In addition to galectin-3, the autophagy adapter protein P62 is also recruited to intracellular Mtb in an ESX-1-dependent manner.^15,21^ We observed that neither prazosin nor gefitinib restored recruitment of P62 to intracellular ESX-1 mutant Mtb to levels comparable to wild-type Mtb (Figure 4 A-B). We therefore hypothesized that, in addition to membrane disruption, autophagy could have a role in mediating cross-presentation of Mtb antigens following ESX-1-mediated phagosome membrane disruption, perhaps by changing the composition of the compartment in which Mtb resides to facilitate antigen processing and loading on MHC-I.^44^ To test this hypothesis, we quantified Mtb antigen presentation on MHC-I in THP-1 macrophages in which ATG7 had been knocked out. ATG7 is required for Mtb to be targeted by the autophagy pathway,^45^ and ATG7 knockout THP-1 macrophages showed impaired clearance of P62-positive autophagic cargo (Supplementary Figure 4). ATG7 knockout THP-1 macrophages infected with Mtb were able to present EsxA_28-36_ and EsxGS_3-11_ on MHC-I, showing that autophagy is dispensable for this process (Figure 4C-D).

**Figure 4.**
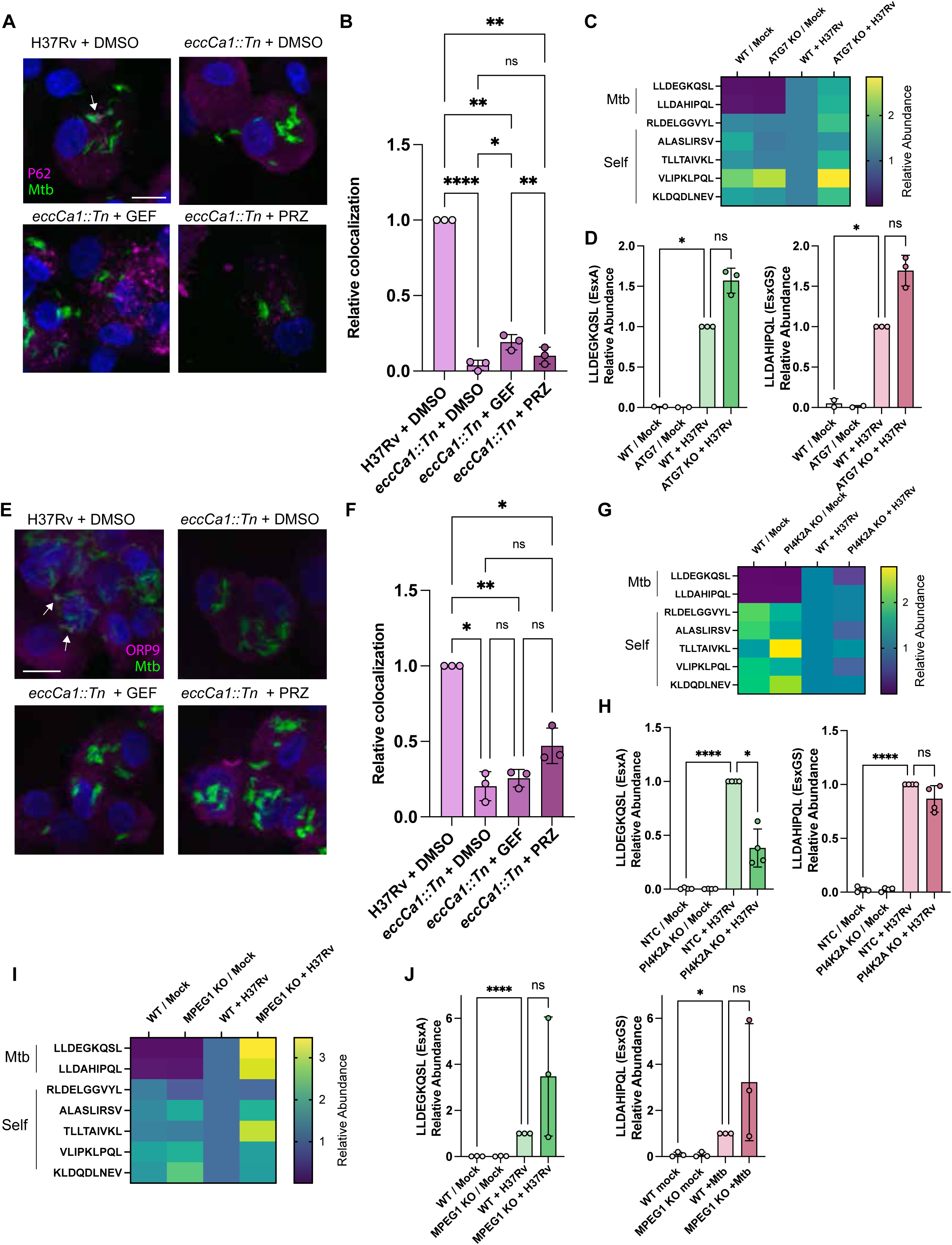
Autophagy, PI4P signaling, and pore formation by perforin-2 (MPEG1) are dispensable for *Mtb* antigen presentation on MHC-I. (A-B) Primary hMDMs infected with wild-type or *eccCa1::Tn Mtb* at MOI 2.5 for 72 hours, treated with prazosin, gefitinib, or DMSO alone for 2 hours, and stained for P62. A) Representative fluorescence confocal microscope images. White arrows indicate intracellular bacteria colocalized with P62. B) Relative quantification of colocalization between intracellular *Mtb* and P62, as measured by analysis of local fluorescence intensity correlation (see Methods) for n = 3 donors (* p < 0.05, ** p < 0.01, **** p < 0.0001, one-way ANOVA with Tukey’s multiple comparisons test). (C-D) ATG7 knockout THP-1 macrophages or the parental wild-type line were infected with *Mtb* at MOI 2.5 and harvested for MHC-I peptide quantification by SureQuant 24 hours post-infection. C) Heatmap of relative abundances of MHC-I peptides (mean of n = 3 replicate experiments). D) Relative abundance of EsxA_28-36_ and EsxGS_3-11_ presented on MHC-I for n = 3 replicate experiments. (E-F) Primary hMDMs infected with wild-type or *eccCa1::Tn Mtb* at MOI 2.5 for 72 hours, treated with prazosin, gefitinib, or DMSO alone for 2 hours, and stained for ORP9. E) Representative fluorescence confocal microscope images. White arrows indicate intracellular bacteria colocalized with ORP9. F) Relative quantification of colocalization between intracellular *Mtb* and ORP9, as measured by analysis of local fluorescence intensity correlation (see Methods) for n = 3 donors (* p < 0.05, ** p < 0.01, one-way ANOVA with Tukey’s multiple comparisons test). (G-H) PI4K2A knockout THP-1 macrophages or the parental wild-type line were infected with *Mtb* at MOI 2.5 and harvested for MHC-I peptide quantification by SureQuant 24 hours post-infection. G) Heatmap of relative abundances of MHC-I peptides (mean of n = 3 replicate experiments). H) Relative abundance of EsxA_28-36_ and EsxGS_3-11_ presented on MHC-I for n = 3 replicate experiments (* p < 0.05, one-way paired ANOVA with Dunnett’s multiple comparisons test). (I-J) MPEG1 knockout THP-1 macrophages or the parental wild-type line were infected with *Mtb* at MOI 2.5 and harvested for MHC-I peptide quantification by SureQuant 24 hours post-infection. I) Heatmap of relative abundances of MHC-I peptides (mean of n = 3 replicate experiments). J) Relative abundance of EsxA_28-36_ and EsxGS_3-11_ presented on MHC-I for n = 3 replicate experiments.

### Endomembrane damage repair mediated by PI4K2A is not strictly required for Mtb-derived antigen presentation on MHC-I

Another process that occurs downstream of endomembrane compartment damage is the formation of contacts between the endoplasmic reticulum (ER) and the damaged compartment(s).^46^ When endomembrane compartments such as lysosomes are ruptured, PI4K2A catalyzes production of phosphoinositide-4-phosphate (PI4P), which ORP9 and ORP11 bind as part of a complex that includes VAPA and VAPB, bridging the ER membrane and that of the damaged endomembrane compartment to facilitate membrane repair.^46^ PI4K2A was also a statistically significant hit in a genetic screen for factors required for cross-presentation in dendritic cells,^3^ which would be consistent with a role for PI4P signaling in facilitating presentation of antigens derived from phagosomal cargo on MHC-I. We therefore asked whether ESX-1-mediated and/or drug-mediated phagosome membrane disruption induces ER-phagosome contacts. Like P62, the ER-phagosome membrane contact site marker ORP9 is recruited to Mtb-containing phagosomes in an ESX-1-dependent manner, and its co-localization with ESX-1 mutant Mtb is not restored to wild-type levels upon treatment with prazosin or gefitinib (Figure 4E-F).

To test whether PI4P signaling and/or ER-phagosome membrane contacts are required for presentation of Mtb antigens on MHC-I, we made a THP-1 cell line in which PI4K2A was knocked out (Supplementary Figure 5) and quantified presentation of EsxA_28-36_ and EsxGS_3-11_ on MHC-I via SureQuant. Presentation of EsxA_28-36_ was reduced in PI4K2A knockout cells compared to wild-type cells, whereas presentation of EsxGS_3-11_ was not affected (Figure 4G-H). These results suggest that ER-phagosome membrane contacts induced following ESX-1-mediated membrane disruption may have a previously unappreciated effect on the efficiency with which certain Mtb antigens can be loaded on MHC-I, although these contacts are apparently not strictly required for presentation of Mtb antigens.

We had initially hypothesized that ER-phagosome membrane contacts might be required for Mtb MHC-I antigen presentation and that prazosin and gefinitinib might fail to rescue presentation of EsxGS_3-11_ despite inducing phagosome membrane damage because of their inability to recapitulate these ER-phagosome membrane contacts. However, knocking out PI4K2A only affected presentation of EsxA_28-36_ and not EsxGS_3-11_, suggesting this hypothesis is not the case. Nonetheless, these results suggest that ER-phagosome membrane contacts can quantitatively influence the efficiency with which some Mtb antigens are presented.

### Perforin-2 (MPEG1) is not required for Mtb-derived antigen presentation on MHC-I

Our prior results showing ESX-1-dependent presentation of Mtb MHC-I antigens are consistent with a model in which antigen translocation to the host cytosol requires a process driven by Mtb itself rather than solely by host cross-presentation pathways.^15^ Here, we sought to test the alternative hypothesis that antigen translocation could instead be mediated by host cell cross-presentation mechanisms, perhaps recruited as a downstream consequence of ESX-1 activity. A CRISPR-Cas9 genetic screen performed by Rodríguez-Silvestre et al.^3^ previously identified macrophage expressed gene 1 (MPEG1) as a pore-forming protein (subsequently termed Perforin-2) that facilitates translocation of proteins from endosomal or phagosomal cargo to the cytosol to enable entry into MHC-I antigen processing pathways. To test whether permeabilization of phagosomes by MPEG1 could be involved in presentation of Mtb antigens, we infected MPEG1 knockout THP-1 macrophages with Mtb and quantified presentation of EsxA_28-36_ and EsxGS_3-11_ by targeted MS. These Mtb MHC-I antigens were presented at comparable or higher levels in *MPEG1* knockout cells compared to wild-type, showing that MPEG1 is dispensable for presentation of Mtb antigens on MHC-I (Figure 4H-I). These results are consistent with the hypothesis that Mtb itself drives translocation of T7SS secreted antigens to the cytosol in a manner dependent on ESX-1 activity, rather than host cross-presentation pathways mediating this translocation.

### Mtb mutants reveal distinct requirements for membrane disruption and MHC-I antigen presentation

To further test whether host cross-presentation pathways are involved in presentation of Mtb MHC-I antigens, we sought to determine whether other Mtb antigens besides the epitopes specifically targeted in our SureQuant analyses could be presented in an ESX-1-independent manner – that is, potentially via host-driven cross-presentation rather than ESX-1 activity. We performed untargeted MS analyses of the MHC-I repertoire of hMDMs infected with ESX-1 mutant Mtb, using two techniques that improve the depth of coverage with which of pathogen-derived MHC-I peptides can be detected (Figure 5A): 1) offline fractionation of peptides prior to MS analysis^15^, and 2) PathMHC, a workflow that uses computational analyses of untargeted MS data to identify infection-specific peaks and target those peaks for more sensitive and selective identification in a follow-up targeted analysis.^37^ Using each of these methods, Mtb-derived peptides were detected in the MHC-I repertoire of hMDMs infected with WT Mtb, but not ESX-1 mutant Mtb (Figure 5B). We attempted to validate one putative ID of an MHC-I peptide derived from PPE3 from an immunopeptidome analysis of macrophages infected with ESX-1 mutant Mtb, but this ID could not be confirmed by SureQuant (i.e., the putative biological peptide did not co-elute with the corresponding synthetic standard; Supplementary Figure 6), suggesting it is likely a false positive.

**Figure 5.**
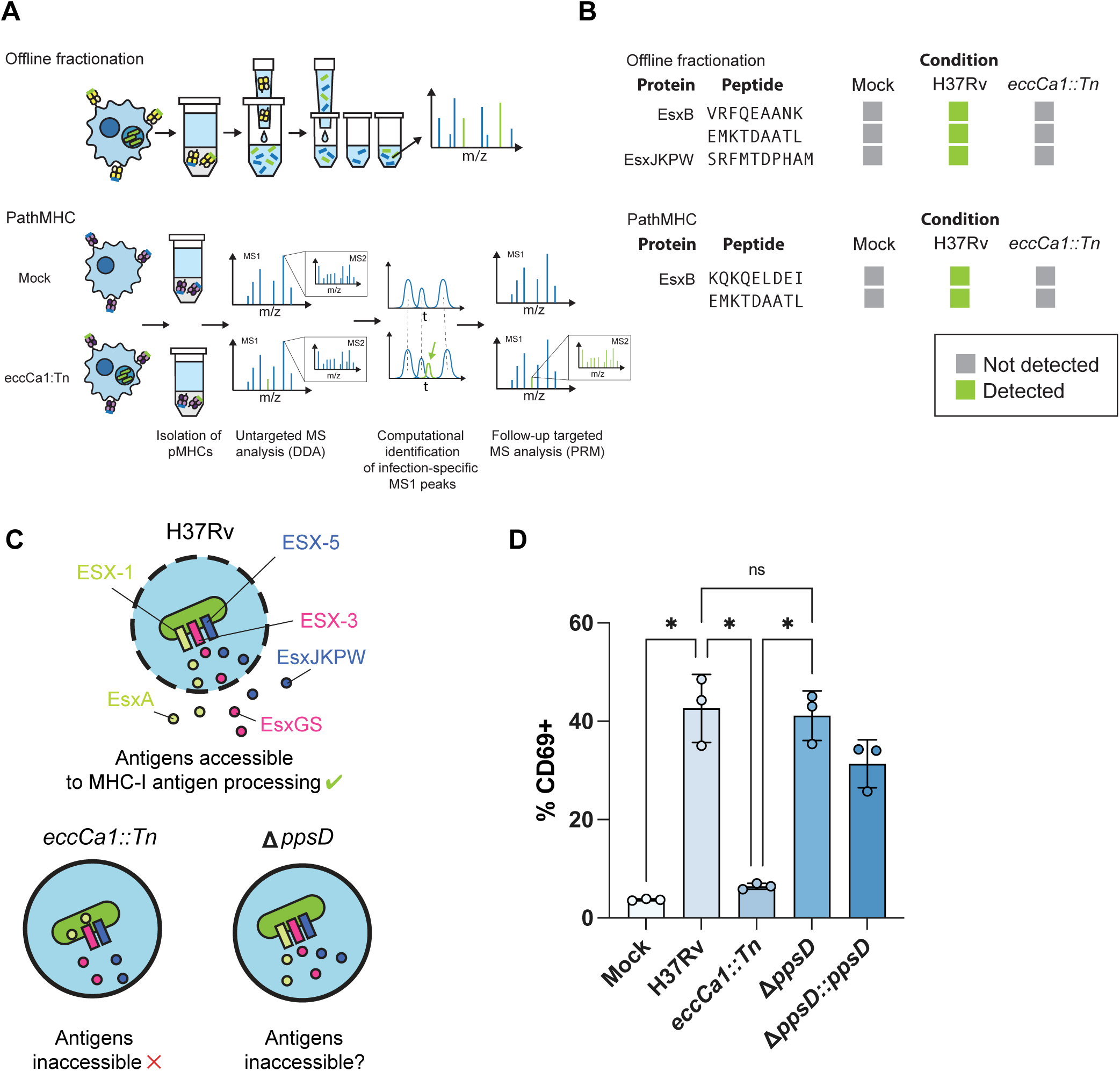
Presentation of Mtb-derived peptides identified by MS requires ESX-1 activity and is not phenocopied by a PDIM mutant. A) Schematic illustration of approaches for in-depth analysis of the MHC-I repertoire of *Mtb*-infected macrophages, including off-line fractionation and PathMHC. B) *Mtb* peptides identified by untargeted analyses of the MHC-I repertoire of hMDMs infected with wild-type or *eccCa1::Tn Mtb*, using offline fractionation or PathMHC. C) Schematic illustrating hypothesized mechanism of antigen release from the phagosome following membrane damage induced by ESX-1 or PDIM. D) Quantification of EsxJKPW_24-34_-specific T cell activation by Mtb-infected macrophages infected with wild-type, *eccCa1::Tn,* Δ*ppsD,* Δ*ppsD::ppsD Mtb*.

Multiple Mtb mutations that disrupt recruitment of galectins to the phagosome have been identified.^47,47–49^ Mtb mutations that disrupt synthesis of the virulence lipid phthiocerol dimycocerosate (PDIM) abrogate type 1 interferon production and galectin recruitment.^25,27,47^ We reasoned that we could leverage Mtb mutants lacking ESX-1 activity or PDIM to disambiguate between a specific role for ESX-1 activity compared to a general role for Mtb-associated phagosomal membrane damage (Figure 5C). We infected hMDMs expressing HLA-B*57:01 with WT Mtb, *eccCa1::Tn*, Δ*ppsD*, or a *ppsD* complemented strain. Given the agreement between our immunopeptidomics results and our co-culture experiments with the EsxJKPW_24-34_-specific reporter T cell line, we used T cell co-culture experiments given the reduced cell requirement of T cell co-culture experiments relative to immunopeptidomics. hMDMs were infected at a multiplicity of infection of 2 and infection was allowed to proceed for 72 hours prior to the addition of cognate SKW3 reporter T cells. After 24 hours of co-culture, we performed flow cytometry to quantify T cell activation, using CD69 as an activation marker. We observed that cells infected with WT Mtb, Δ*ppsD*, or the complemented strain all elicited T cell activation. By contrast, we saw a specific defect in T cell activation exclusively with the ESX-1-deficient mutant (Figure 5D). This result was unexpected because PDIM and ESX-1 are thought to work in concert to enable Mtb access to the host cytosol.^47^ The specific disruption of Mtb-derived antigen presentation by a mutation in ESX-1 but not a mutation in the PDIM biosynthesis pathway is consistent with a model in which the processes that drive galectin recruitment to phagosomes (putatively gross membrane rupture) are dispensable and insufficient for antigen access to MHC-I processing pathways, whereas ESX-1 activity is specifically required. This result is consistent with a model in which ESX-1 facilitates antigen presentation through selective translocation to the cytosol rather than nonspecific membrane rupture, or a general defect in T7SS secretion in intracellular Mtb in the absence of ESX-1 activity.

## Discussion

To detect and eliminate intracellular pathogens, the immune system relies on detection of antigens that become accessible to antigen processing pathways within the host cell and are subsequently presented on MHCs. Antigens contained within a live microbial pathogen may be inaccessible to these processing pathways unless they are secreted or otherwise translocated to a compartment surveilled by host antigen processing machinery. In the case of pathogens that can survive inside phagosomes, the immune system relies in part on cross-presentation pathways in which the host cell translocates antigens from endomembrane spaces to the cytosol for processing and presentation on MHC-I.^50^ However, pathogens themselves also face evolutionary pressure to deliver effector proteins into the host cell to manipulate host cell physiology, access nutrients, or otherwise create a favorable and secure intracellular niche.^51–53^ By using specialized secretion systems to accomplish this, pathogens potentially deliver their own antigens to host antigen processing pathways. Here, we investigated the potential role of the ESX-1 secretion system of Mtb in enabling bacterial antigens to access MHC-I processing pathways.

Quantitative targeted MS provides a means of sensitively and reliably quantifying pathogen-derived MHC epitopes under a range of experimental perturbations to understand how host and pathogen processes contribute to antigen presentation. In this study, we took advantage of this technique to probe the determinants of Mtb antigen presentation on MHC-I in infected macrophages, finding that ESX-1 enables antigen presentation on MHC-I through a cytosolic pathway, in a manner independent of both autophagy and MPEG1-mediated cross-presentation. Drug-induced phagosome membrane disruption could not functionally substitute for ESX-1 activity to mediate MHC-I antigen presentation. Additionally, ER-phagosome membrane contacts were formed in an ESX-1-dependent manner and modulated the efficiency of presentation of an Mtb MHC-I epitope, but were not essential for MHC-I antigen presentation. Whereas cross-presentation pathways in myeloid cells can enable presentation of antigens derived from phagosomal cargo, our results are consistent with a model in which ESX-1 activity enables Mtb antigens to access the host cytosol independently of host-driven cross-presentation pathways, and processed peptides are subsequently imported into the ER for loading on MHC-I.

While transduction of Mtb-infected cells with viral proteins that inhibit TAP has previously been shown to inhibit activation of Mtb-specific CD8+ T cells,^54^ our results showing that presentation of EsxA_28-36_ and EsxGS_3-11_ is abolished in TAP-deficient cells provide additional evidence consistent with the hypothesis that Mtb MHC-I antigens are presented through a cytosolic pathway. The fact that Esx-family Mtb MHC-I antigens access the cytosol may have implications for TB vaccine design. Multiple vaccine platforms based on nucleic acids (mRNA vaccines and viral vectors)^55,56^ have been clinically validated through their successful use against SARS-CoV-2 and now represent potential tools for cytosolic delivery of vaccine antigens in other disease areas. Delivery of Mtb Esx-family vaccine antigens to cytosolic antigen processing pathways may help recapitulate presentation of the same Mtb MHC-I epitopes that are presented during infection. These approaches may help prime CD8+ T cell responses capable of recognizing Mtb-infected cells.

Previous studies have shown that *Mycobacterium marinum* ESX-1 activity upregulates PI4K2A, ORP9, and other members of a membrane repair pathway dependent on ER-phagosome contacts. Additionally, oxysterol binding protein (OSBP) – a PI4P and cholesterol antiporter, also a member of this pathway – is recruited to *M. marinum* phagosomes in an ESX-1-dependent manner.^57^ Here, we similarly show that ORP9 is recruited to Mtb-containing phagosomes in an ESX-1-dependent manner and reveal a possible role for ER-phagosome membrane contacts in facilitating presentation of an Mtb MHC-I epitope (EsxA_28-36_), as knocking out PI4K2A led to less efficient presentation. ER-phagosome membrane contacts could potentially enable more efficient TAP-mediated import of Mtb antigens into the ER for loading on MHC-I by bringing phagosome membranes and ER membrane into proximity. These membrane contacts most likely do not facilitate antigen presentation through direct communication between the ER lumen and phagosome lumen, as this access would presumably be TAP-independent, whereas presentation of EsxA_28-36_ is TAP-dependent. It is currently unclear why knocking out PI4K2A modulates presentation of EsxA_28-36_ but not EsxGS_3-11_. Factors such as differences in the subcellular location where proteolytic processing of the antigen occurs or the susceptibility of the peptide to internal cleavage by cytosolic proteases could contribute to determining which MHC-I peptides are influenced by ER contacts. The potential role of ER contacts with phagosomes and other endomembrane compartments in coordinating loading of antigens on MHC-I warrants further investigation, both in the setting of Mtb infection and in the context of other phagosomal cargoes. However, these contacts are not strictly required for presentation and cannot be the primary mechanism underlying the dependence of MHC-I antigen presentation on ESX-1.

One possible implication of the finding that ESX-1 activity facilitates presentation of many Mtb MHC-I antigens is that mutations that down-regulate ESX-1 activity could hypothetically enable Mtb to evade recognition by CD8+ T cells. Mtb clinical isolates have been reported that are capable of causing disease despite bearing loss-of-function mutations in genes essential for ESX-1 function,^58^ although it is unclear whether these strains gain any fitness advantage through immune evasion that could compensate for the loss of other fitness-conferring functions of ESX-1. Naturally occurring mutations that increase ESX-1 activity have also been reported in clinical isolates.^59^ Examining how variation in levels of ESX-1 activity affects antigen presentation and immune recognition across Mtb genomic diversity could be an important subject for future studies to evaluate the risk that naturally occurring mutations could help Mtb evade vaccines targeting antigens presented in an ESX-1-dependent manner.

Several possible models could explain why drug-induced phagosome membrane disruption does not rescue presentation of Mtb antigens on MHC-I in the absence of ESX-1 activity. ESX-1 activity enables Mtb to manipulate host cells in a variety of ways, including facilitating translocation of T7SS effectors that manipulate endomembrane trafficking and repair throughout the cell,^60^ disruption of mitochondrial function,^41^ and modulation of cell death pathways.^61^ Some of these other functions of ESX-1 or their indirect consequences might also be required to enable Mtb antigen presentation on MHC-I. Another possibility is that rather than releasing proteins from the phagosome non-specifically, the ESX-1 system or one or more of its substrates is involved in creating pores or channels to selectively translocate T7SS substrates to the host cytosol, analogous to the pore-forming complexes inserted into host membranes by type III secretion systems.^62,63^ Treatment with a membrane-disrupting drug may not recapitulate this coordinated translocation process. Alternatively, interdependent regulation of the activity of other T7SSs by ESX-1 or by ESX-1-mediated changes in the phagosomal environment could lead to reduced export of ESX-3 and ESX-5 substrates by intracellular Mtb in the absence of ESX-1 activity, even though these secretion systems operate independently in axenic culture.^42^ In the absence of an assay to directly measure secretion of ESX-3 and ESX-5 substrates in infected cells, this currently remains unclear.

Recruitment of membrane damage markers such as galectins to phagosomes has often been measured as a proxy for rupture of the phagosome membrane,^64^ which would potentially enable antigen access to the cytosol. However, we show here that a mutation in the PDIM biosynthesis pathway, a process previously shown to be required for recruitment of membrane damage markers to Mtb phagosomes,^47^ does not abrogate antigen presentation to a T cell line specific for an Mtb MHC-I epitope. This result implies that antigen access to cytosolic MHC-I processing pathways and recruitment of marker proteins typically associated with membrane rupture may be two separable processes. We and others in the field had previously speculated that the known role of ESX-1 in rupture of the phagosome membrane could be a driver of antigen access to MHC-I processing pathways,^15,65^ but our results here suggest that recruitment of membrane damage markers like galectins to Mtb phagosomes is separable from access to MHC-I processing pathways. Other prior studies showed that PDIM export is required for recruitment of galectins to phagosomes containing intracellular *Mycobacterium marinum*, but not for hemolytic activity, whereas ESX-1 is required for both.^66^ This finding suggests that ESX-1 systems may disrupt host membranes in multiple distinct ways, some of which could be involved in antigen translocation but not galectin recruitment and vice versa.

Lewinsohn et al. previously showed that a CD8+ T cell clone specific for TB8.4, a protein containing a Sec pathway secretion signal peptide, recognizes Mtb-infected dendritic cells in an ESX-1-independent manner.^54^ Our results in this study show that ESX-1 activity is required for presentation of an important class of antigens (Esx-family proteins) that our prior results show are prominently represented in the MHC-I repertoire and are commonly targeted by vaccine candidates, but this ESX-1-dependence may not hold true for all antigens in all infected cell types. Other studies have also found that priming of CD8+ T cells specific for EsxH (an ESX-3 substrate) does not differ between infection with wild-type and *espA* knockout Mtb *in vivo*.^67^ As EspA is required for secretion of EsxA and EsxB,^38,68^ and the latter are in turn required for recruitment of membrane damage and autophagy markers to phagosomes,^21^ this result has been interpreted to mean that ESX-1 activity is dispensable for *in vivo* CD8+ T cell priming against substrates of other secretion systems. If this is the case, it would imply that – unlike *in vitro* macrophages – some types of infected antigen presenting cells are able to present Mtb antigens on MHC-I independently of ESX-1 activity, or that EsxH can be encountered and cross-presented by cells that are not directly infected. However, our results showing that membrane damage marker recruitment and antigen access to MHC-I are not always coterminous suggests a second possible explanation – namely that knocking out *espA* ablates the ability of ESX-1 to drive membrane damage marker recruitment, but not the function(s) required for MHC-I antigen presentation.

Besides Mtb, other intracellular bacterial pathogens possess virulence mechanisms that disrupt or translocate across phagosome membranes, enabling bacterial antigens to access cytosolic MHC-I antigen processing pathways. The type III secretion systems of bacterial pathogens such as *Salmonella* and *Yersinia* sp. can translocate proteins to the host cytosol that are then presented on MHC-I,^14,69^ and *Listeria monocytogenes* secretes pore-forming proteins that rupture the phagosome to provide access to the host cytosol.^13^ Our results here suggest that the ESX-1 system of Mtb may serve as an analogous mechanism of pathogen-driven access to cytosolic antigen processing pathways. Previous preclinical studies have exploited the *Salmonella* type III secretion system to deliver cancer antigens to MHC-I processing pathways,^69–71^ and mycobacterial systems capable of enhancing antigen presentation on MHC-I could hypothetically be put to similar uses. BCG is already used therapeutically in the treatment of bladder cancer^72^ and has previously been engineered to express the ESX-1 system of *Mycobacterium marinum*, enhancing CD8+ T cell responses to multiple mycobacterial antigens.^73^ The ESX-1 system could potentially facilitate delivery of recombinantly expressed antigens as well, if this can be accomplished without increasing the virulence of attenuated mycobacteria like BCG. Other bacterial secretion systems also translocate effector proteins to the host cytosol in order to manipulate host cell physiology, including the type IV secretion systems of *Legionella pneumophila* and *Coxiella burnetii*.^74,75^ A broad range of intracellular pathogens may therefore face a tradeoff between the benefits of using their secretion systems to access and manipulate the host cell and the potential liability of detection by CD8+ T cells through antigen presentation on MHC-I.

Our results in this study suggest that host cross-presentation pathways alone do not mediate efficient access of Mtb Esx-family antigens to MHC-I processing pathways. Rather, the activity of a bacterial secretion system is required to facilitate presentation, a distinct mechanism from cross-presentation of many other phagosomal cargoes. Our results emphasize the distinction between viral or cytosol-resident pathogens on the one hand, with abundant access to cytosolic host MHC-I processing pathways, and pathogens that reside in endomembrane compartments on the other, for which additional mechanisms are required for pathogen-derived proteins to access host MHC-I antigen processing pathways. We add to a body of evidence consistent with a model in which Mtb antigens are presented via access to the host cytosol and subsequent import into the ER for loading on MHC-I. We also identify a possible role for ER-phagosome membrane contacts in coordinating the import of Mtb antigens released from phagosomes into the ER for loading on MHC-I. These results advance the field’s understanding of the cell biology of antigen presentation in Mtb infection and may help inform delivery of Mtb vaccine antigens.

## Limitations of the study

Mtb-derived peptides presented on MHC-I and MHC-II are rare compared to self peptides due to the replication rate of the bacterium and the low biomass of Mtb proteins accessible to host antigen processing machinery compared to host proteins. As a consequence, we only have a limited number of peptides we can track for a given HLA allotype. There are hundreds of known HLA allotypes, and we have not studied these dynamics across all HLA allotypes. We are currently unable to track most Mtb antigens with monoclonal antibodies or label specific Mtb antigens with epitope tags to track their localization in human macrophages, so we use TAP-dependent presentation as an indirect measure for access of these antigens to the cytosol. Mutations in *eccCa1*, *ppsD*, and other ESX-1- and PDIM-related genes have previously been associated with defects in intracellular growth and survival of *Mtb*^24,25,68,76,77^ that are difficult to fully decouple from antigen production, secretion, and presentation. Although we and others^78^ have included controls for bacterial burden when comparing antigen presentation from WT and ESX-1-deficient *Mtb*, impairments in intracellular bacterial growth or metabolism could still affect antigen presentation in a manner not directly related to secretion or cytosolic translocation. Lastly, there are several Mtb lineages with slight differences in their behavior in macrophages. We used H37Rv as our strain of choice because we have useful Mtb mutants in this genetic background, but the generalizability of our conclusions to all Mtb isolates remains to be determined.

## Methods

### Statistical tests

Unless otherwise noted, all statistical tests were performed using GraphPad Prism 10.

#### Generation of a natively paired D504 TCRα:β amplicon

Total RNA was extracted from the D504 T cell clone and natively paired TCRα:β cDNA was generated using our established two-step, semi-nested PCR workflow to maintain α–β pairing. Briefly, an initial semi-nested suppression PCR incorporating blocking oligonucleotides complementary to the 3′ ends of unfused TCRα and TCRβ products was performed using a HotStart GoTaq Polymerase System (Promega, Madison, WI). A second semi-nested PCR was performed using a KAPA HiFi HotStart PCR Kit (Roche, Basel, Switzerland). PCR products were resolved by agarose gel electrophoresis and purified from 1.5% SYBR Safe agarose (Thermo Fisher Scientific, Waltham, MA).

#### Vector construction to express the D504 TCR variable regions

Purified D504 TCRα:β amplicons and a modified pLVX-EF1α-IRES-mCherry vector (Takara Bio, Mountain View, CA) were digested with BstBI and AgeI and gel-purified. Inserts were ligated into the vector with T4 DNA Ligase (New England Biolabs, Ipswich, MA), cleaned with a DNA Clean & Concentrator kit (Zymo Research, Irvine, CA), and transformed by electroporation into MegaX DH10B T1 Electrocomp Cells (Thermo Fisher Scientific). Plasmids were prepared using a ZymoPURE II Maxiprep kit (Zymo Research). The remaining expression cassette—comprising the distal portion of the TCRβ constant region, a P2A self-cleaving peptide, and a modified TCRα leader sequence engineered to contain an MluI site—was then introduced between the TCRβ and TCRα variable regions using SpeI and MluI (New England Biolabs) to enable co-expression of the heterodimeric D504 TCR.

Following transformation, colonies were screened by Sanger sequencing using vector-complementary primers EF1-α (forward, 5′-TCAAGCCTCAGACAGTGGTTC-3′) and TRAC M-Inner (reverse, 5′-GCTCTTGAAGTCCATAGACCTCA-3′). CE-derived TCR variable sequences were queried by IgBLAST to identify functional in-frame rearrangements. A single clone (colony #15) contained functional in-frame TRAV and TRBV sequences for D504 and was advanced. Whole-plasmid sequencing (Plasmidsaurus) confirmed the results. IgBLAST comparison indicated two FR1-region deviations in TRBV relative to germline (one nonsynonymous, one synonymous). Single-nucleotide corrections were introduced into the TRBV FR1 region by site-directed mutagenesis on the D504 plasmid backbone (D504_F9_pLVX). The FR1-corrected TRBV construct was subsequently used for downstream experiments.

#### Lentiviral packaging and transduction of SKW3 cells

Lentivirus was produced by transfecting adherent 293FT cells (Thermo Fisher Scientific) at 70–90% confluence in T75 flasks with 12 µg pLVX-EF1α-IRES-mCherry-D504, 8 µg psPAX2, and 2 µg pMD2.G (Addgene, Watertown, MA) using Lipofectamine 3000 with P3000 reagent in Opti-MEM (Thermo Fisher Scientific) per flask. After 72 h at 37 °C, supernatants were clarified (800 × g, 10 min) and concentrated with Lenti-X Concentrator (Takara Bio) overnight at 4 °C, then pelleted (1,500 × g, 45 min, 4 °C) and resuspended in RPMI 1640. For transduction, 3 × 10^6^ SKW3 cells (DSMZ, Braunschweig, Germany) were plated in 3 mL RPMI 1640 in a 6-well plate and infected with 400 µL thawed viral stock in the presence of 4 µg/mL polybrene (Sigma-Aldrich, St. Louis, MO) overnight at 37 °C with 5% CO2. Cells were washed, transferred to T25 flasks in pre-warmed RPMI 1640, expanded for 3 days, then washed with PBS and sorted by flow cytometry for internal mCherry expression to enrich D504 TCR-expressing SKW3 cells.

#### Primary human cell isolation, differentiation, and culture

Deidentified buffy coats were obtained from Massachusetts General Hospital. Samples are acquired and provided to research groups with no identifying information. HLA-A*02:01+, HLA-B*57:01+ leukapheresis samples (leukopaks) were obtained from StemCell. PBMCs were isolated by density-based centrifugation using Ficoll (GE Healthcare). CD14+ monocytes were isolated from PBMCs using a CD14 positive-selection kit (Stemcell). Isolated monocytes were differentiated in phenol red free R10 media [RPMI 1640 without phenol red (Gibco) supplemented with 10% heat-inactivated FBS (Gibco), 1% HEPES (Corning), 1% L-glutamine (Sigma)] supplemented with 25 ng/mL M-CSF (Biolegend, 572902). Monocytes were cultured on appropriate ultra-low-attachment plates or flasks (Corning) for 6 days to differentiate into macrophages.

#### THP-1 macrophage culture and differentiation

THP-1 cells were cultured in R10 media [RPMI 1640 (Gibco) supplemented with 10% heat-inactivated FBS (Gibco), 1% HEPES (Corning), 1% L-glutamine (Sigma)] and grown to a density of 1 million cells per mL in T25 flasks, then differentiated into macrophages via treatment for 24 hours with 150 nM phorbol myristate acetate followed by 24 hours in fresh media without drug.

#### Mtb culture

*Mycobacterium tuberculosis* (Mtb) H37Rv was grown in Difco Middlebrook 7H9 media supplemented with 10% OADC, 0.2% glycerol, and 0.05% Tween-80 to mid-log phase. Strains expressing episomal GFP were grown in media supplemented with 50 μg/mL hygromycin B (Sigma-Aldrich).

#### Mtb infection of macrophages

The Mtb culture was pelleted by centrifugation, washed once with PBS, resuspended in R10 media and centrifuged at low speed (500 rpm for 5 minutes) to pellet clumps, leaving a uniform suspension of bacteria in the supernatant. Macrophages were infected at the indicated multiplicity of infection (MOI) for 4 hr and then washed with PBS to remove extracellular Mtb. Infected macrophages were cultured in R10 media for the remainder of the experiment.

#### Intracellular bacterial RNA sequencing

Infected bacterial RNA sequencing was performed as previously described.^79^ Briefly, monocytes were isolated and differentiated as described above, then replated at 1 million per condition in a 6-well ultra-low attachment plate. Macrophages were infected at MOI 3 then washed after 4 hours. After 24 hours, cells were lysed in Trizol, and intact bacteria were lysed by bead beating. RNA was isolated using chloroform and processed for sequencing using the Agilent SureSelect XT HS2 RNA Kit and custom probes complementary to the H37Rv transcriptome to enrich for bacterial transcripts.

#### hipMHC internal standard preparation

UV-mediated peptide exchange and quantification of hipMHC complexes by ELISA were performed as previously described.^80^ Amidated peptides with sequences AL[+7]ADGVQKV-NH2, ALNEQIARL[+7]-NH2, and SLPEEIGHL[+7]-NH2 were used to make hipMHC standards.

#### MHC immunoprecipitation

Macrophages were harvested by collecting the culture media (containing any cells in suspension), washing adherent cells with PBS, incubating remaining adherent cells with PBS supplemented with 4 mM EDTA for 15 minutes at 37 °C, gently scraping with a cell scraper, collecting the detached cells, and washing the flask with PBS and collecting the wash. Harvested cells were pelleted by centrifugation. For optimization experiments, 25 million Raji cells, which grow in suspension, were collected by centrifugation without additional treatment. The harvested cells were then washed with PBS, and lysed in 1 mL of MHC lysis buffer [20 mM Tris, 150 mM sodium chloride, pH 8.0, supplemented with 1% CHAPS, 1 x HALT protease and phosphatase inhibitor cocktail (Pierce), and 0.2 mM phenylmethylsulfonyl fluoride (Sigma-Aldrich)].

Lysate from Mtb-infected or mock-infected cells was sonicated using a Q500 ultrasonic bath sonicator (Qsonica) in five 30-second pulses at an amplitude of 60%, cleared by centrifugation at 16,000 x g for 5 minutes, and sterile filtered twice using 0.2 μm filter cartridges (Pall NanoSep).

For quantitative SureQuant analyses, the protein concentration of the lysates was normalized by BCA assay (Pierce), and 100 fmol of each hipMHC standard was added to each sample. Lysates were then added to protein A sepharose beads pre-conjugated to pan-MHC-I antibody (clone W6/32), prepared as previously described,^81^ and incubated rotating at 4 °C overnight (12–14 hours). Beads were then washed and peptide-MHC complexes eluted as previously described.^81^

#### MHC-I peptide isolation

MHC-I-associated peptides were purified using 10 kDa molecular weight cutoff filters (Pall NanoSep) as previously described,^81^ snap-frozen in liquid nitrogen, and lyophilized.

#### Offline fractionation

Where offline fractionation is indicated, lyophilized peptides were resuspended in solvent A2 (10 mM triethyl ammonium bicarbonate [TEAB], pH 8.0) and loaded onto a fractionation column (a 200 μm inner diameter fused silica capillary packed in-house with 10 cm of 10 μM C18 beads). Peptides were fractionated using an Agilent 1100 series liquid chromatograph using buffers A2 (10 mM TEAB, pH 8.0) and B2 (99% acetonitrile, 10 mM TEAB, pH 8.0). The fractionation column was washed with solvent A2, and peptides were separated using a gradient of 1–5% solvent B2 over 5 min, 5–40% over 60 min, 40–70% over 10 min, hold for 9 min, and 70%–1% over 1 min. Ninety-second fractions were collected, concatenated into 12 tubes, over 90 min in total. Fractions were flash-frozen in liquid nitrogen and lyophilized.

#### DDA MS analyses

MHC-I peptide samples were resuspended in 0.1% formic acid. 25% of the sample was refrozen and reserved for later SureQuant validation analyses, while 75% of the sample was used for DDA analysis. For all MS analyses, samples were analyzed using an Orbitrap Exploris 480 mass spectrometer (Thermo Fisher Scientific) coupled with an UltiMate 3000 RSLC Nano LC system (Dionex), Nanospray Flex ion source (Thermo Fisher Scientific), and column oven heater (Sonation). The MHC peptide sample was loaded onto a fused silica capillary chromatography column with an integrated electrospray tip (∼1 μm orifice) prepared and packed in-house with 10 cm of 1.9 μm C18 beads (ReproSil-Pur).

Standard mass spectrometry parameters were as follows: spray voltage, 2.0 kV; no sheath or auxiliary gas flow; ion transfer tube temperature, 275 °C. The Orbitrap Exploris 480 mass spectrometer was operated in data dependent acquisition (DDA) mode. Peptides were eluted using a gradient of 6–25% buffer B (70% Acetonitrile, 0.1% formic acid) over 75 min, 25–45% over 5 min, 45–100% over 5 min, hold for 1 min, and 100% to 3% over 2 min. Full scan mass spectra (350–1200 m/z, 60,000 resolution) were detected in the orbitrap analyzer after accumulation of 3×10^6^ ions (normalized AGC target of 300%) or 25ms. For every full scan, MS/MS scans were collected during a 3 s cycle time. Ions were isolated (0.4 m/z isolation width) using the standard AGC target and automatic determination of maximum injection time, fragmented by HCD with 30% CE, and scanned at a resolution of 120,000. Charge states <2 and >4 were excluded, and precursors were excluded from selection for 30 s if fragmented n=2 times within a 20-s window.

#### Synthetic standard survey MS analyses

Synthetic SIL standard peptides were synthesized by BioSynth as a crude peptide library, with the exception of SIL EsxJKPW_24-34_, which was synthesized by the Koch Institute biopolymers core facility as described previously.^15^ DDA MS analysis of the SIL peptide mixture was performed as described above (see DDA MS analyses) with the following modifications: Peptides were eluted using a flow rate of 300 nL/min and a gradient of 6–35% buffer B over 30 min, 35–45% over 2 min, 45–100% over 3 min, and 100% to 2% over 1 min. No dynamic exclusion was used.

A second set of survey analyses was performed on the mixture of SIL peptides with background matrix using the full SureQuant acquisition method (see below). SIL peptides were spiked into a sample of MHC-I peptides purified as described above from THP-1 cells, providing a representative background matrix. Because SIL amino acids are not 100% pure, SIL peptide concentrations were adjusted and survey analyses were repeated until the SIL peptide could be reliably detected while minimizing background signal detected at the mass of the biological peptide.

#### PathMHC analyses

For PathMHC analyses, PRM inclusion lists were generated as previously described.^37^ In PRM analyses, inclusion list masses were matched by m/z and charge state, with a mass tolerance of 5 ppm. Acquisition was scheduled for a window of 3 minutes centered on the observed retention time of the target precursor ion in the prior DDA analysis in which it was originally detected. Precursors were excluded from selection for 4 seconds after being fragmented 1 time.

#### SureQuant analyses

Standard MS parameters and MS1 scan parameters were as described above (see DDA MS analyses). The custom SureQuant acquisition template available in Thermo Orbitrap Exploris Series 2.0 was used for method building. For each method, the optimal charge state and most intense product ions were determined via a survey analysis of the synthetic SIL peptide standards. One method branch was created for each m/z offset between the SIL peptide and biological peptide as previously described.^15^ A threshold of n=3 out the top 6 product ions was used for pseudo-spectral matching, with a mass accuracy tolerance of 10 ppm. As none of the targets of these analyses has an m/z < 380, an MS1 scan range of m/z 380–1500 was used to exclude some common background ions.

Targeted MS data were analyzed using Skyline Daily Build 22.1.9.208. For each target peptide, the intensities of three product ions with the highest ratio of true signal in a positive sample to background in a negative control were integrated over the time during which the peptide was scanned. These intensities were normalized by the integrated intensities of the corresponding product ions from the corresponding SIL standard over the same time interval, and these ratios were averaged. Finally, these averaged ratios were normalized by the average ratio of the integrated intensities of the hipMHC standard peptides and normalized to the reference condition for a given experiment (typically wild-type or untreated macrophages infected with Mtb) to obtain the final relative abundance of each target. 1 pmol of SIL EsxJKPW_24-34_ and 100 fmol of each other SIL trigger peptide were spiked into each sample per analysis.

#### DDA data search and manual inspection

All mass spectra were analyzed with Proteome Discoverer (PD, version 3.0) and searched using Sequest with rescoring using INFERYS and Percolator against a custom database comprising the Uniprot human proteome (UP000005640) together with the Uniprot *Mycobacterium tuberculosis* H37Rv proteome (UP000001584). No enzyme was used, and variable modifications included oxidized methionine for all analyses. Peptide-spectrum matches from MHC-I analyses were filtered with the following criteria: search engine rank = 1, length between 8 and 12 amino acids, XCorr ≥ 2.0, spectral angle ≥ 0.6, and percolator q-value < 0.05.

Identifications (IDs) of putative Mtb-derived peptides were rejected if any peptide-spectrum matches (PSMs) for the same peptide were found in the unfiltered DDA MS data for the corresponding mock-infected control. For each putative Mtb peptide identified, MS/MS spectra and extracted ion chromatograms (XIC) were manually inspected, and the ID was only accepted for further validation if it met the following criteria: (1) MS/MS spectra contained enough product ions separated by the mass of a single amino acid to unambiguously assign a majority of the peptide sequence; (2) neutral losses were consistent with the chemical properties of the peptide; (3) manual *de novo* sequencing did not reveal an alternate peptide sequence that would explain a greater number of MS/MS spectrum peaks; (4) XIC showed a peak in MS intensity at the mass to charge ratio (m/z) of the peptide precursor ion at the retention time at which it was identified that did not appear in the corresponding mock-infected control.

#### Immunofluorescence microscopy

100,000 macrophages per well plated on chamber slides (Ibidi) were infected with GFP-expressing Mtb at MOI 2.5 or mock-infected with media containing no Mtb. At the indicated time point post-infection, macrophages were washed with PBS and fixed with 4% paraformaldehyde (PFA) in PBS for 1 hr. After the full time course was collected, all slides were blocked with 5% normal goat serum in PBS supplemented with 0.3% v/v Triton X-100 for 1 hr at room temperature and stained overnight at 4°C with primary antibody ([Ab clones]) diluted in antibody dilution buffer (PBS with 1% w/v bovine serum albumin and 0.3% v/v triton X-100). Wells were washed three times with PBS for 5 min each and stained with Alexa fluor 647 (AF647)-conjugated goat anti-rabbit or anti-mouse secondary antibody (Thermo) diluted to a final concentration of 1 μg/mL in antibody dilution buffer at room temperature for 2 hr. Wells were washed three times with PBS for 5 min each and stained with DAPI at a final concentration of 300 nM in PBS for 15 min at room temperature. Wells were washed three times with PBS for 5 min each, well dividers were removed, and slide covers were mounted on slides using Prolong Diamond anti-fade mounting media (Thermo). Slides were imaged using a TissueFAXS Confocal slide scanner system (TissueGnostics). 25 fields of view were acquired per condition using a 40x objective lens.

#### Segmentation of Mtb-containing phagosomes

Image segmentation was performed using opencv-python 4.5.4.60. A two-dimensional Gaussian blur with a standard deviation of five pixels was applied to de-noise GFP fluorescence images. A binary mask was generated from the blurred GFP image using a fluorescence intensity threshold that was empirically selected for each biological replicate. GFP+ phagosomes were further segmented using the watershed algorithm.

#### Colocalization analysis

A correlation image was generated by computing the Pearson correlation coefficient between the GFP fluorescent intensity and AF647 fluorescent intensity in a 41×41 pixel sliding window with a step size of 1. This correlation value was averaged over each Mtb-containing phagosome, and phagosomes with a mean value of greater than or equal to 0.6 were considered co-localized. Code can be found at https://github.com/oleddy/local_correlation_analysis.

#### Knockout cell lines

An ATG7 knockout THP-1 cell line and corresponding parental wild-type control were obtained from Abcam (ab277836). An MPEG1 knockout cell line and corresponding parental wild-type control were obtained from Synthego (custom order). A PI4K2A knockout THP-1 cell line was generated by nucleofecting THP-1 cells with ribonucleoprotein complexes comprising Cas9 and single guide RNAs (sgRNAs), following the approach of Jani et al.^49^ 180 pmol of sgRNA (Synthego) and 20 pmol of purified Cas9 protein (Synthego) were diluted to a final volume of 25 μL in Nucleofector Solution with the provided supplement (Lonza). After incubating for 10 minutes, the RNP complexes were added to 2 ✕ 10^5^ THP-1 cells and nucleofected using the SG Cell Line 4D-Nucleofector X Kit S (Lonza) following the manufacturer instructions.

#### Isolation of clonal THP-1 knockout cell lines

Following nucleofection with Cas9-RNPs, clonal THP-1 lines were isolated by limiting dilution. Cells were diluted to a density of 1 cell per 400 μL in sterile-filtered conditioned media from wild-type THP-1 cultures, and 200 μL were added to each well of a 96-well plate. Clones were allowed to expand for 5 weeks and screened for PI4K2A protein production by western blot (see below).

#### Generation of TAP1-knockout THP-1 cells

THP-1 cells (ATCC) were transduced with VSV-G-pseudotyped lentivirus encoding lentiCas9-Blast. After 3 days, transduced cells were selected in culture media containing 5 μg/mL blasticidin (Thermo Fisher) for >7 days to generate Cas9^+^ THP-1 cells. Cas9^+^ THP-1 cells were then subsequently transduced with VSV-G-pseudotyped lentivirus encoding a lentiGuide-Puro vector containing an sgRNA targeting *TAP1* (5’-TCGAAGCTTTGCCAACGAGG-3’). After 72 h, transduced cells were selected in culture media containing 1 μg/mL puromycin (Thermo Fisher) for >7 days. Puromycin-selected cells were then stained with PE-conjugated pan-MHC-I antibody (clone W6/32) and sorted for TAP1 knockout (MHC-I^low^) cells using the Sony SH800 Cell Sorter. Sorted TAP1-knockout THP-1 cells were subsequently expanded and validated for homogeneous loss of TAP1 (i.e., low MHC-I expression) via flow cytometry staining before using for downstream assays. The lentiCas9-Blast and lentiGuide-Puro plasmids used for lentiviral transduction were gifts from Feng Zhang (Addgene plasmids # 52962 and 52963).^82^

#### Western blot

Undifferentiated THP-1 cell clones isolated as described above following nucleofection with Cas9-RNPs were grown to a density of 1 million cells per well in 6-well tissue culture plates. Cells were washed with PBS and lysed in high-SDS lysis buffer [1% SDS in 50 mM Tris-HCl (pH 8.0) with 1 mM PMSF, and HALT protease inhibitors (ThermoFisher, 78429)]. Lysates were incubated at 95° C for 5 minutes, then diluted with 1.25x RIPA without SDS (50 mM Tris, 187.5 mM NaCl, 0.625% sodium deoxycholate, 1.25% Triton X-100) to achieve a final concentration of 1X RIPA lysis buffer with 0.2% SDS. Lysates were incubated at room temperature for 5 minutes, then cleared by centrifugation at 10,000 ✕ g for 10 minutes at 4° C. Protein concentration was normalized by absorbance at 660 nm. NuPAGE LDS Sample Buffer (Thermo) and NuPAGE Sample Reducing Agent (Thermo) were added to a final concentration of 1x, and 20 μL of lysate per well was separated by SDS-PAGE. Protein was transferred to a PVDF membrane using an iBlot2 dry transfer system (Thermo). The PVDF membrane was stained with Ponceau S reagent (ThermoFisher, A40000279) for 5 minutes at room temperature, washed 2x with water for 60 seconds and imaged using a GelDoc (BioRad). Membranes were blocked for 1 hr at room temperature with blocking buffer (Licor) and incubated overnight at 4 °C with anti-PI4K2A antibody (Santa Cruz Biotechnology, sc-390026), diluted 1:1000 in antibody diluent (Licor). Blots were washed three times for 5 minutes each with tris-buffered saline with 0.1% tween-20 (TBS-T) and incubated for 1 hr at room temperature with IRDye 800CW goat anti-rabbit secondary antibody (Licor) and IRDye 680CW goat anti-mouse secondary antibody (Licor), each diluted 1:1000 in antibody diluent (Licor). Blots were washed three times for five minutes each with TBS-T and imaged using an Odyssey DLx imaging system (Licor).

## Acknowledgements

The authors thank all the staff members of the Ragon Institute, the Koch Institute, and MIT for the essential work they do to make our research possible. We thank Lauren Stopfer, Cameron Flower, Tigist Tamir, Elizabeth Choe, Ryuhjin Ahn, Cal Gunnarsson, Christine Zheng, Charul Jani, and Amy Barczak for helpful conversations, training, and technical guidance. The EsxJ-specific T cell clone was generously provided by Deborah and David Lewinsohn. The *eccCa1::Tn* Mtb strain and corresponding parental wild-type control were generously provided by Amy Barczak and Charul Jani. This work is supported by funding from the MIT Center for Precision Cancer Research at the Koch Institute, the Koch Institute Frontier Research Program, American Cancer Society grant RSG-22-134-01-IBCD, U.S. Department of Defense grant W81XWH1810296, and U.S. NIH grants U01 CA238720, U54 CA283114, 1R35GM142900, and R01A1022553. O.L. was supported in part by U.S. NIEHS grant T32-ES007020. This work was performed in part in the Ragon Institute BSL3 core facility, which is supported by the NIH-funded Harvard University Center for AIDS Research (P30 AI060354). We thank Yong Xie, Julie Boucau, and Eliane Shwairi for managing the facility.

## Figure captions

**Supplementary Figure 1. Loss of ESX-1 activity disrupts activation of an EsxJ-specific T cell reporter in a TCR-dependent manner**

hMDMs expressing HLA-B*57:01 underwent a mock infection or infection with wild-type Mtb H37Rv or *eccCa1::Tn* (ESX-1 mutant) Mtb at MOI 2.5 for 72 hours prior to coculture with SKW3 reporter T cells expressing a TCR specific for EsxJKPW_24-34_ or HIV Gag_22-30_ (RM9) at a ratio of 1 million T cells to 1 million hMDMs for 24 hours.

**Supplementary Figure 2. Prazosin treatment does not rescue presentation of *Mtb* peptides by THP-1 macrophages**

THP-1 macrophages were infected with wild-type or *eccCa1::Tn Mtb* at MOI 2.5 for 24 hours, treated with prazosin or DMSO alone for 2 hours, and harvested for MHC-I antigen quantification by SureQuant.

A) Heatmap of relative abundances of MHC-I peptides (mean of n = 2 replicate experiments).

B) Relative abundance of EsxA_28-36_ presented on MHC-I for n = 2 replicate experiments.

C) Relative abundance of EsxGS_3-11_ presented on MHC-I for n = 2 replicate experiments.

**Supplementary Figure 3. Bacterial burden is only modestly reduced in macrophages infected with ESX-1-deficient *Mtb* relative to wild-type**

Quantification of GFP+ area in infected hMDMs quantified by fluorescence microscopy 72 hours post-infection for n = 6 donors, normalized by DAPI+ area to account for host cell density. All differences not statistically significant (one-way ANOVA with Tukey’s multiple comparisons test).

**Supplementary figure 4. ATG7 knockout THP-1 macrophages accumulate P62+ autophagic cargo**

Representative fluorescence confocal microscopy images of wild-type or ATG7 knockout THP-1 macrophages stained for P62 by immunofluorescence, 24 hours post-infection with *Mtb* at MOI 2.5.

**Supplementary Figure 5. Validation of a PI4K2A knockout THP-1 cell line made using Cas9 ribonucleoprotein (RNP) complex nucleofection**

PI4K2A western blot comparing single THP-1 clones isolated from populations nucleofected with Cas9 RNPs containing a PI4K2A-specific pool of single guide RNAs (sgRNAs) or a non-targeting control (NTC) pool of sgRNAs.

**Supplementary Figure 6. A putative ID of an MHC-I peptide derived from PPE3 did not pass validation using SureQuant**

Fragment ion chromatograms for PPE3_42-50_ (sequence VADELIGLL) from SureQuant analyses of MHC-I peptides from *Mtb*-infected or uninfected cells (sample reserved from initial untargeted discovery analysis).

**Supplementary Table 1. Relative transcript abundance in wild-type and ESX-1 mutant Mtb**

## References

1. Gonzales, G. A. et al. The pore-forming apolipoprotein APOL7C drives phagosomal rupture and antigen cross-presentation by dendritic cells. Sci. Immunol. (2024).

2. Canton, J. et al. The receptor DNGR-1 signals for phagosomal rupture to promote cross-presentation of dead-cell-associated antigens. Nat. Immunol. 22, 140–153 (2021).

3. Rodríguez-Silvestre, P. et al. Perforin-2 is a pore-forming effector of endocytic escape in cross-presenting dendritic cells.

4. Chen, C. Y. et al. A Critical Role for CD8 T Cells in a Nonhuman Primate Model of Tuberculosis. PLoS Pathog. 5, e1000392 (2009).

5. Flynn, J. L., Goldstein, M. M., Triebold, K. J., Koller, B. & Bloom, B. R. Major histocompatibility complex class I-restricted T cells are required for resistance to Mycobacterium tuberculosis infection. Proc. Natl. Acad. Sci. 89, 12013–12017 (1992).

6. Patankar, Y. R. et al. Limited recognition of Mycobacterium tuberculosis-infected macrophages by polyclonal CD4 and CD8 T cells from the lungs of infected mice. Mucosal Immunol. 13, 140–148 (2020).

7. Nyendak, M. et al. Adenovirally-Induced Polyfunctional T Cells Do Not Necessarily Recognize the Infected Target: Lessons from a Phase I Trial of the AERAS-402 Vaccine. Sci. Rep. 6, 36355 (2016).

8. Blanchard, N. et al. Immunodominant, protective response to the parasite Toxoplasma gondii requires antigen processing in the endoplasmic reticulum. Nat. Immunol. 9, 937–944 (2008).

9. Mantegazza, A. R., Magalhaes, J. G., Amigorena, S. & Marks, M. S. Presentation of Phagocytosed Antigens by MHC Class I and II. Traffic 14, 135–152 (2013).

10. Cruz, F. M., Colbert, J. D., Merino, E., Kriegsman, B. A. & Rock, K. L. The Biology and Underlying Mechanisms of Cross-Presentation of Exogenous Antigens on MHC-I Molecules. Annu. Rev. Immunol. 35, 149–176 (2017).

11. Song, R. & Harding, C. V. Roles of proteasomes, transporter for antigen presentation (TAP), and beta 2-microglobulin in the processing of bacterial or particulate antigens via an alternate class I MHC processing pathway. J. Immunol. 156, 4182–4190 (1996).

12. Gonzales, G. A. et al. The pore-forming apolipoprotein APOL7C drives phagosomal rupture and antigen cross-presentation by dendritic cells. Sci. Immunol. (2024).

13. Brunt, L. M., Portnoy, D. A. & Unanue, E. R. Presentation of Listeria monocytogenes to CD8+ T cells requires secretion of hemolysin and intracellular bacterial growth. J. Immunol. 145, 3540–3546 (1990).

14. Rüssmann, H. et al. Yersinia enterocolitica-mediated translocation of defined fusion proteins to the cytosol of mammalian cells results in peptide-specific MHC class I-restricted antigen presentation. Eur. J. Immunol. 30, 1375–1384 (2000).

15. Leddy, O., White, F. M. & Bryson, B. D. Immunopeptidomics reveals determinants of Mycobacterium tuberculosis antigen presentation on MHC class I. eLife 12, e84070 (2023).

16. Stanley, S. A., Raghavan, S., Hwang, W. W. & Cox, J. S. Acute infection and macrophage subversion by Mycobacterium tuberculosis require a specialized secretion system. Proc. Natl. Acad. Sci. 100, 13001–13006 (2003).

17. Siegrist, M. S. et al. Mycobacterial Esx-3 is required for mycobactin-mediated iron acquisition. Proc. Natl. Acad. Sci. 106, 18792–18797 (2009).

18. Ates, L. S. et al. Essential Role of the ESX-5 Secretion System in Outer Membrane Permeability of Pathogenic Mycobacteria. PLOS Genet. 11, e1005190 (2015).

19. Mott, D. et al. High Bacillary Burden and the ESX-1 Type VII Secretion System Promote MHC Class I Presentation by *Mycobacterium tuberculosis* –Infected Macrophages to CD8 T Cells. J. Immunol. 210, 1531–1542 (2023).

20. Simeone, R. et al. Phagosomal Rupture by Mycobacterium tuberculosis Results in Toxicity and Host Cell Death. PLoS Pathog. 8, e1002507 (2012).

21. Watson, R. O., Manzanillo, P. S. & Cox, J. S. Extracellular M. tuberculosis DNA Targets Bacteria for Autophagy by Activating the Host DNA-Sensing Pathway. Cell 150, 803–815 (2012).

22. Grotzke, J. E., Siler, A. C., Lewinsohn, D. A. & Lewinsohn, D. M. Secreted Immunodominant *Mycobacterium tuberculosis* Antigens Are Processed by the Cytosolic Pathway. J. Immunol. 185, 4336–4343 (2010).

23. Lewinsohn, D. M. et al. Human Mycobacterium tuberculosis CD8 T Cell Antigens/Epitopes Identified by a Proteomic Peptide Library. PLoS ONE 8, e67016 (2013).

24. Guinn, K. M. et al. Individual RD1□region genes are required for export of ESAT□6/CFP□10 and for virulence of *Mycobacterium tuberculosis*. Mol. Microbiol. 51, 359–370 (2004).

25. Lerner, T. R. et al. Phthiocerol dimycocerosates promote access to the cytosol and intracellular burden of Mycobacterium tuberculosis in lymphatic endothelial cells. BMC Biol. 16, 1 (2018).

26. Bell, S. L., Lopez, K. L., Cox, J. S., Patrick, K. L. & Watson, R. O. Galectin-8 Senses Phagosomal Damage and Recruits Selective Autophagy Adapter TAX1BP1 To Control *Mycobacterium tuberculosis* Infection in Macrophages. mBio 12, e01871–20 (2021).

27. Barczak, A. K. et al. Systematic, multiparametric analysis of Mycobacterium tuberculosis intracellular infection offers insight into coordinated virulence. PLOS Pathog. 13, e1006363 (2017).

28. Manzanillo, P. S., Shiloh, M. U., Portnoy, D. A. & Cox, J. S. Mycobacterium Tuberculosis Activates the DNA-Dependent Cytosolic Surveillance Pathway within Macrophages. Cell Host Microbe 11, 469–480 (2012).

29. Stanley, S. A., Johndrow, J. E., Manzanillo, P. & Cox, J. S. The Type I IFN Response to Infection with *Mycobacterium tuberculosis* Requires ESX-1-Mediated Secretion and Contributes to Pathogenesis. J. Immunol. 178, 3143–3152 (2007).

30. MacGurn, J. A. & Cox, J. S. A Genetic Screen for *Mycobacterium tuberculosis* Mutants Defective for Phagosome Maturation Arrest Identifies Components of the ESX-1 Secretion System. Infect. Immun. 75, 2668–2678 (2007).

31. Love, M. I., Huber, W. & Anders, S. Moderated estimation of fold change and dispersion for RNA-seq data with DESeq2. Genome Biol. 15, 550 (2014).

32. Zhu, A., Ibrahim, J. G. & Love, M. I. Heavy-tailed prior distributions for sequence count data: removing the noise and preserving large differences. Bioinformatics 35, 2084–2092 (2019).

33. Blum, J. S., Wearsch, P. A. & Cresswell, P. Pathways of Antigen Processing. Annu. Rev. Immunol. 31, 443–473 (2013).

34. Leddy, O. et al. Validation and quantification of peptide antigens presented on MHCs using SureQuant. Nat. Protoc. 10.1038/s41596-024-01076-x (2024) doi:10.1038/s41596-024-01076-x.

35. Axelsson-Robertson, R. et al. A Broad Profile of Co-Dominant Epitopes Shapes the Peripheral Mycobacterium tuberculosis Specific CD8+ T-Cell Immune Response in South African Patients with Active Tuberculosis. PLoS ONE 8, e58309 (2013).

36. Lewinsohn, D. A. et al. Immunodominant Tuberculosis CD8 Antigens Preferentially Restricted by HLA-B. PLoS Pathog. 3, e127 (2007).

37. Leddy, O., Yuki, Y., Carrington, M., Bryson, B. D. & White, F. M. PathMHC: a workflow to selectively target pathogen-derived MHC peptides in discovery immunopeptidomics experiments for vaccine target identification.Preprint at 10.1101/2024.09.11.612454 (2024).

38. Fortune, S. M. et al. Mutually dependent secretion of proteins required for mycobacterial virulence. Proc. Natl. Acad. Sci. 102, 10676–10681 (2005).

39. Cronin, R. M., Ferrell, M. J., Cahir, C. W., Champion, M. M. & Champion, P. A. Proteo-genetic analysis reveals clear hierarchy of ESX-1 secretion in *Mycobacterium marinum*. Proc. Natl. Acad. Sci. 119, e2123100119 (2022).

40. Kozik, P. et al. Small Molecule Enhancers of Endosome-to-Cytosol Import Augment Anti-tumor Immunity. Cell Rep. 32, 107905 (2020).

41. Pagán, A. J. et al. mTOR-regulated mitochondrial metabolism limits mycobacterium-induced cytotoxicity. Cell 185, 3720–3738.e13 (2022).

42. Champion, P. A. D., Stanley, S. A., Champion, M. M., Brown, E. J. & Cox, J. S. C-Terminal Signal Sequence Promotes Virulence Factor Secretion in *Mycobacterium tuberculosis*. Science 313, 1632–1636 (2006).

43. Lewinsohn, D. M. et al. Human Mycobacterium tuberculosis CD8 T Cell Antigens/Epitopes Identified by a Proteomic Peptide Library. PLoS ONE 8, e67016 (2013).

44. English, L. et al. Autophagy enhances the presentation of endogenous viral antigens on MHC class I molecules during HSV-1 infection. Nat. Immunol. 10, 480–487 (2009).

45. Aylan, B. et al. ATG7 and ATG14 restrict cytosolic and phagosomal Mycobacterium tuberculosis replication in human macrophages. Nat. Microbiol. 8, 803–818 (2023).

46. Tan, J. X. & Finkel, T. A phosphoinositide signalling pathway mediates rapid lysosomal repair. Nature 609, 815–821 (2022).

47. Augenstreich, J. et al. ESX-1 and phthiocerol dimycocerosates of *Mycobacterium tuberculosis* act in concert to cause phagosomal rupture and host cell apoptosis. Cell. Microbiol. 19, e12726 (2017).

48. Beckwith, K. S. et al. Plasma membrane damage causes NLRP3 activation and pyroptosis during Mycobacterium tuberculosis infection. Nat. Commun. 11, 2270 (2020).

49. Jani, C. et al. VPS18 contributes to phagosome membrane integrity in Mycobacterium tuberculosis–infected macrophages. Sci. Adv. (2025).

50. Cruz, F. M., Colbert, J. D., Merino, E., Kriegsman, B. A. & Rock, K. L. The Biology and Underlying Mechanisms of Cross-Presentation of Exogenous Antigens on MHC-I Molecules. Annu. Rev. Immunol. 35, 149–176 (2017).

51. Ham, H., Sreelatha, A. & Orth, K. Manipulation of host membranes by bacterial effectors. Nat. Rev. Microbiol. 9, 635–646 (2011).

52. Dean, P. Functional domains and motifs of bacterial type III effector proteins and their roles in infection. FEMS Microbiol. Rev. 35, 1100–1125 (2011).

53. Cascales, E. & Christie, P. J. The versatile bacterial type IV secretion systems. Nat. Rev. Microbiol. 1, 137–149 (2003).

54. Lewinsohn, D. M. et al. Secreted Proteins from *Mycobacterium tuberculosis* Gain Access to the Cytosolic MHC Class-I Antigen-Processing Pathway. J. Immunol. 177, 437–442 (2006).

55. Polack, F. P. et al. Safety and Efficacy of the BNT162b2 mRNA Covid-19 Vaccine. N. Engl. J. Med. 383, 2603–2615 (2020).

56. Corchado-Garcia, J. et al. Analysis of the Effectiveness of the Ad26.COV2.S Adenoviral Vector Vaccine for Preventing COVID-19. JAMA Netw. Open 4, e2132540 (2021).

57. Anand, A. et al. ER-dependent membrane repair of mycobacteria-induced vacuole damage. mBio 14, e00943–23 (2023).

58. Clemmensen, H. S. et al. An attenuated Mycobacterium tuberculosis clinical strain with a defect in ESX-1 secretion induces minimal host immune responses and pathology. Sci. Rep. 7, 46666 (2017).

59. Luna, M. J. et al. Frequently arising ESX-1-associated phase variants influence *Mycobacterium tuberculosis* fitness in the presence of host and antibiotic pressures. mBio 16, e03762–24 (2025).

60. Mittal, E. et al. Mycobacterium tuberculosis Type VII Secretion System Effectors Differentially Impact the ESCRT Endomembrane Damage Response. mBio 9, e01765–18 (2018).

61. Dallenga, T. et al. M. tuberculosis-Induced Necrosis of Infected Neutrophils Promotes Bacterial Growth Following Phagocytosis by Macrophages. Cell Host Microbe 22, 519–530.e3 (2017).

62. Ghosh, P. Process of Protein Transport by the Type III Secretion System. Microbiol. Mol. Biol. Rev. 68, 771–795 (2004).

63. Matteï, P. et al. Membrane targeting and pore formation by the type III secretion system translocon. FEBS J. 278, 414–426 (2011).

64. Liu, F.-T. & Stowell, S. R. The role of galectins in immunity and infection. Nat. Rev. Immunol. 23, 479–494 (2023).

65. Harriff, M. J., Purdy, G. E. & Lewinsohn, D. M. Escape from the Phagosome: The Explanation for MHC-I Processing of Mycobacterial Antigens? Front. Immunol. 3, (2012).

66. Osman, M. M., Pagán, A. J., Shanahan, J. K. & Ramakrishnan, L. Mycobacterium marinum phthiocerol dimycocerosates enhance macrophage phagosomal permeabilization and membrane damage. PLOS ONE 15, e0233252 (2020).

67. Woodworth, J. S., Fortune, S. M. & Behar, S. M. Bacterial Protein Secretion Is Required for Priming of CD8^+^ T Cells Specific for the *Mycobacterium tuberculosis* Antigen CFP10. Infect. Immun. 76, 4199–4205 (2008).

68. Garces, A. et al. EspA Acts as a Critical Mediator of ESX1-Dependent Virulence in Mycobacterium tuberculosis by Affecting Bacterial Cell Wall Integrity. PLoS Pathog. 6, e1000957 (2010).

69. Hegazy, W. A. H., Xu, X., Metelitsa, L. & Hensel, M. Evaluation of Salmonella enterica Type III Secretion System Effector Proteins as Carriers for Heterologous Vaccine Antigens. Infect. Immun. 80, 1193–1202 (2012).

70. Nishikawa, H. In vivo antigen delivery by aSalmonella typhimurium type III secretion system for therapeutic cancer vaccines. J. Clin. Invest. 116, 1946–1954 (2006).

71. Xu, X. et al. Effective Cancer Vaccine Platform Based on Attenuated *Salmonella* and a Type III Secretion System. Cancer Res. 74, 6260–6270 (2014).

72. Alexandroff, A. B., Jackson, A. M., O’Donnell, M. A. & James, K. BCG immunotherapy of bladder cancer: 20 years on. The Lancet 353, 1689–1694 (1999).

73. Gröschel, M. I. et al. Recombinant BCG Expressing ESX-1 of Mycobacterium marinum Combines Low Virulence with Cytosolic Immune Signaling and Improved TB Protection. Cell Rep. 18, 2752–2765 (2017).

74. Hubber, A. & Roy, C. R. Modulation of Host Cell Function by *Legionella pneumophila* Type IV Effectors. Annu. Rev. Cell Dev. Biol. 26, 261–283 (2010).

75. Segal, G., Feldman, M. & Zusman, T. The Icm/Dot type-IV secretion systems of *Legionella pneumophila* and *Coxiella burnetii*. FEMS Microbiol. Rev. 29, 65–81 (2005).

76. Mittal, E. et al. Mycobacterium tuberculosis virulence lipid PDIM inhibits autophagy in mice. Nat. Microbiol. 9, 2970–2984 (2024).

77. Stanley, S. A., Raghavan, S., Hwang, W. W. & Cox, J. S. Acute infection and macrophage subversion by Mycobacterium tuberculosis require a specialized secretion system. Proc. Natl. Acad. Sci. 100, 13001–13006 (2003).

78. Mott, D. et al. High Bacillary Burden and the ESX-1 Type VII Secretion System Promote MHC Class I Presentation by *Mycobacterium tuberculosis* –Infected Macrophages to CD8 T Cells. J. Immunol. 210, 1531–1542 (2023).

79. Gunnarsson, C., McGinn, R. A., Hochfelder, J. & Bryson, B. D. Systems-level characterization of EGFR kinase inhibitors reveals heterogeneous effects on Mtb-macrophage interactions.

80. Stopfer, L. E., Mesfin, J. M., Joughin, B. A., Lauffenburger, D. A. & White, F. M. Multiplexed relative and absolute quantitative immunopeptidomics reveals MHC I repertoire alterations induced by CDK4/6 inhibition. Nat. Commun. 11, 2760 (2020).

81. Stopfer, L. E., Mesfin, J. M., Joughin, B. A., Lauffenburger, D. A. & White, F. M. Multiplexed relative and absolute quantitative immunopeptidomics reveals MHC I repertoire alterations induced by CDK4/6 inhibition. Nat. Commun. 11, 2760 (2020).

82. Sanjana, N. E., Shalem, O. & Zhang, F. Improved vectors and genome-wide libraries for CRISPR screening. Nat. Methods 11, 783–784 (2014).

