## Supplementary figures and images for "Control of antigen presentation on MHC-I by a bacterial secretion system"

### Supplemental Figure 1

**Supplementary Figure 1**

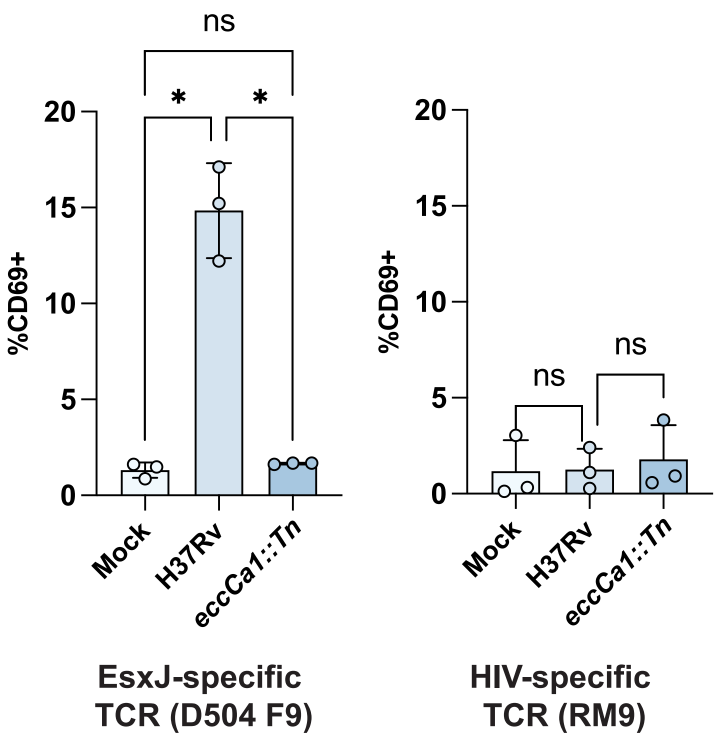

### Supplemental Figure 2

Supplementary Figure 2

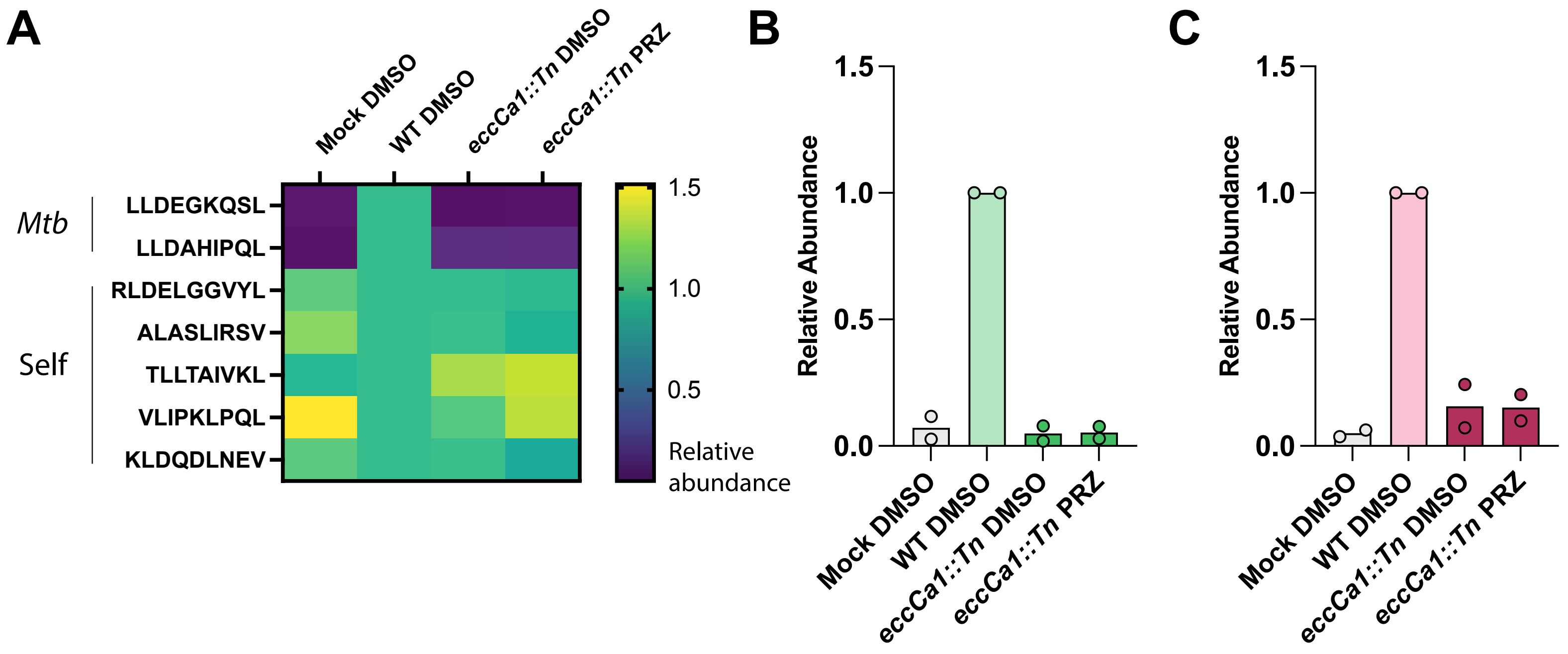

### Supplemental Figure 3

#### Supplemental Figure 3

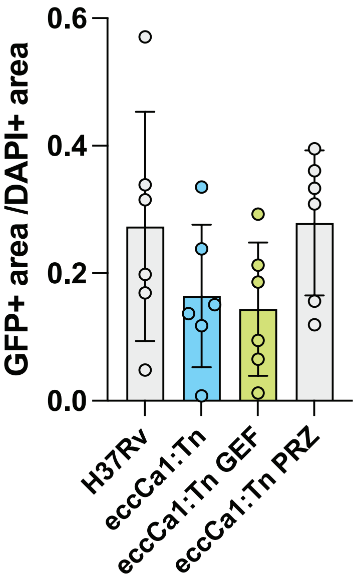

### Supplemental Figure 4

##### Supplementary Figure 4

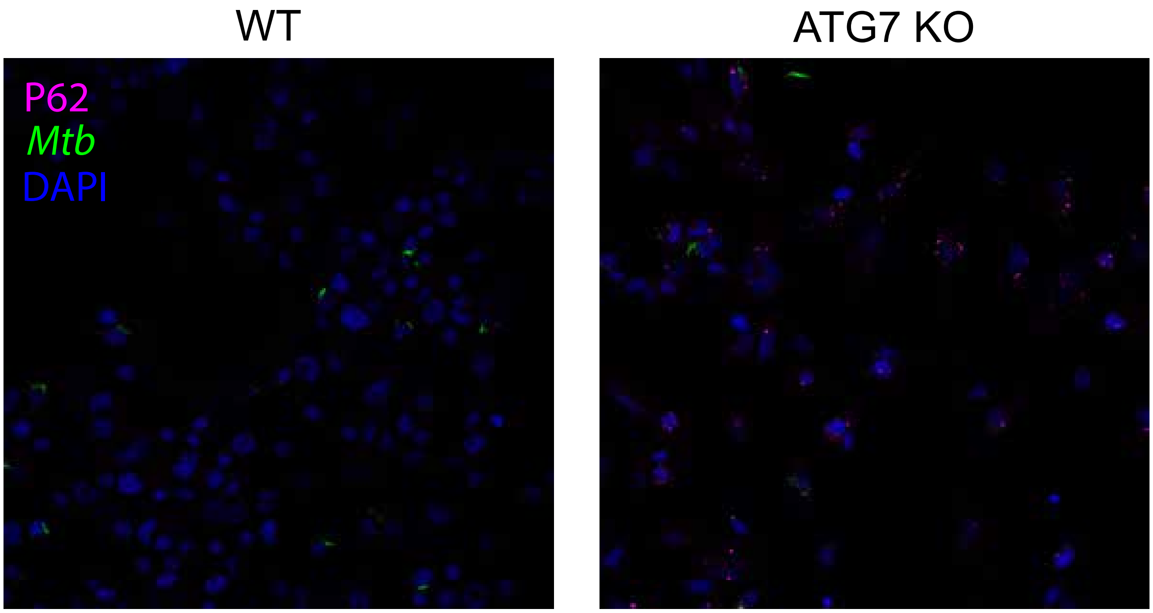

### Supplemental Figure 5

**Supplementary Figure 5**

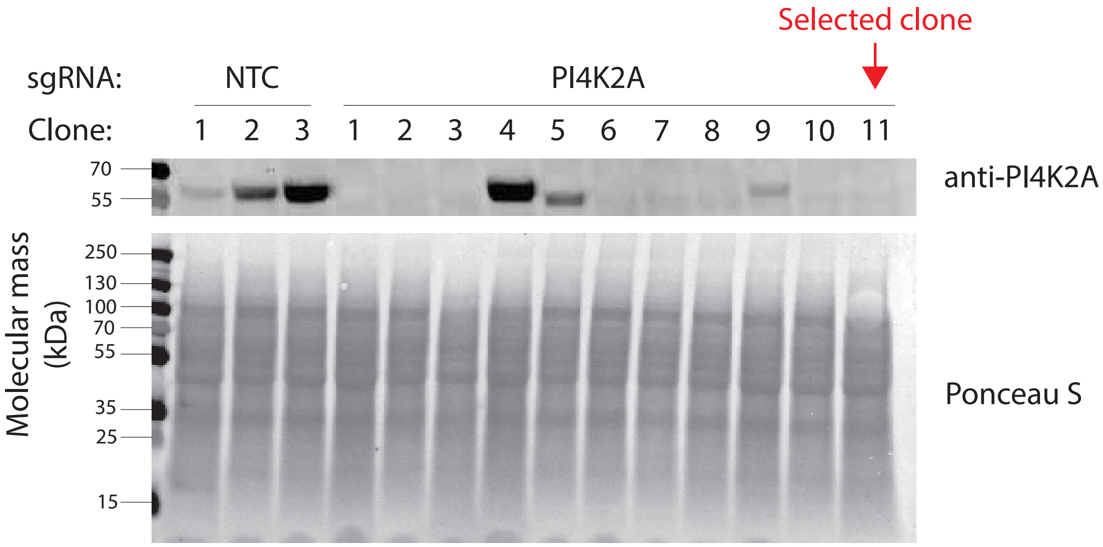

### Supplemental Figure 6

**Supplementary Figure 6**

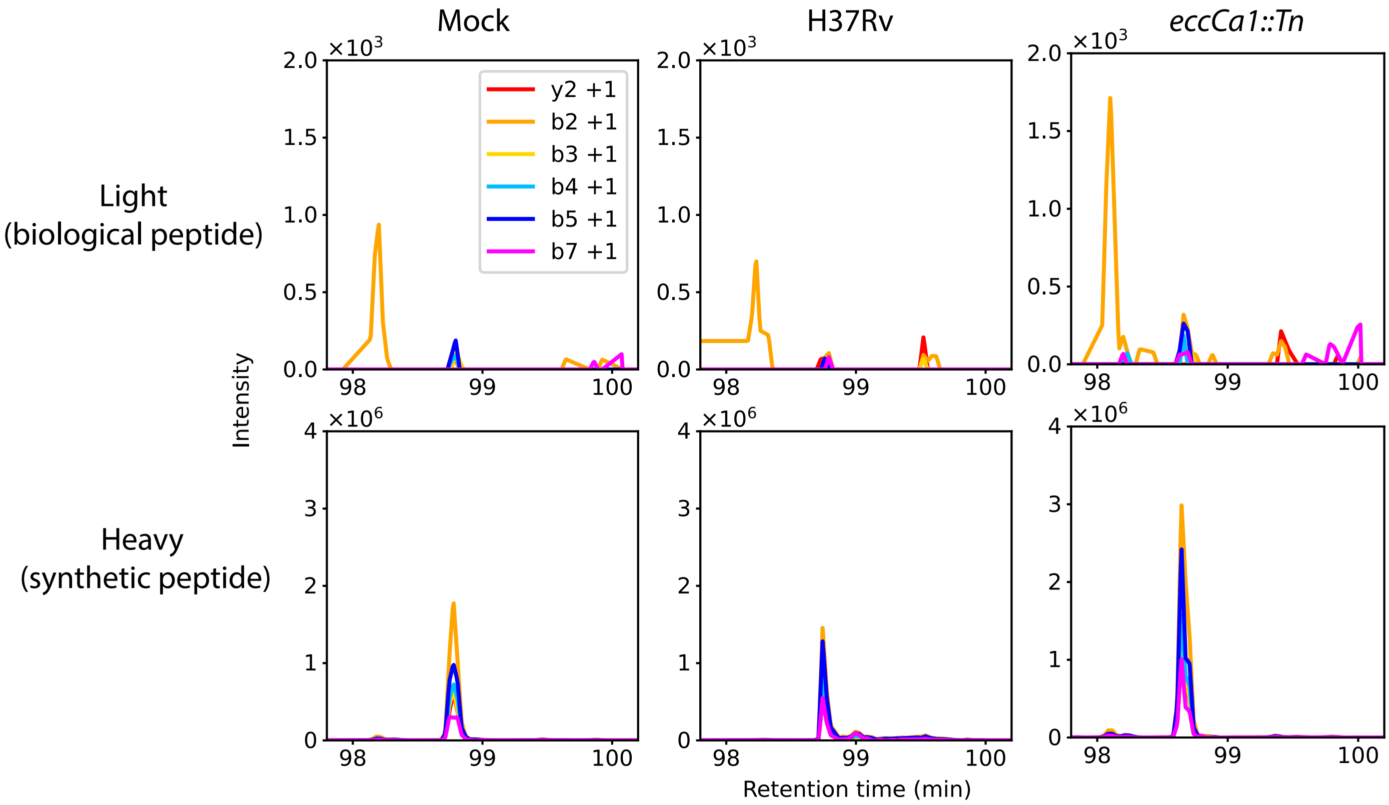
